# Sparsification of AP firing in adult-born hippocampal granule cells via voltage-dependent α5-GABA_A_ receptors

**DOI:** 10.1101/2020.07.22.216200

**Authors:** Meredith Lodge, Maria-Clemencia Hernandez, Jan M. Schulz, Josef Bischofberger

**Author notes:** These authors jointly supervised the work (Jan M. Schulz, Josef Bischofberger). Address correspondence to: Dr. Josef Bischofberger, Department of Biomedicine, University of Basel, Pestalozzistr. 20, CH-4056 Basel, Switzerland.

## Abstract

GABA can depolarize immature neurons close to the action potential (AP) threshold in development and adult neurogenesis. Nevertheless, GABAergic synapses effectively inhibit AP firing in newborn granule cells of the adult hippocampus as early as 2 weeks post mitosis. The underlying mechanisms are largely unclear. Here we analyzed GABAergic inputs in newborn 2- to 4-week-old hippocampal granule cells mediated by soma-targeting parvalbumin (PV) and dendrite-targeting somatostatin (SOM) interneurons. Surprisingly, both interneuron subtypes activate α5-subunit containing GABA_A_ receptors (α5-GABA_A_Rs) in young neurons, showing a nonlinear voltage dependence with increasing conductance around the AP threshold. By contrast, in mature cells, PV interneurons mediate linear GABAergic synaptic currents lacking α5-subunits, while SOM-interneurons continue to target nonlinear α5-GABA_A_Rs. Computational modelling shows that the voltage-dependent amplification of α5-GABA_A_R opening in young neurons is crucial for inhibition of AP firing to generate balanced and sparse firing activity, even with depolarized GABA reversal potentials.

## Introduction

GABA is the major inhibitory transmitter in the adult brain. During embryonic and early postnatal development, however, the GABA_A_-reversal potential is depolarized relative to the resting membrane potential (Ben-Ari, 2002). The depolarizing GABAergic synapses support NMDA receptor (NMDAR) activation and activity-dependent growth of dendrites and synapses, important for proper development of the brain (Wang and Kriegstein, 2008, 2011). Remarkably, depolarizing GABAergic synapses still exert inhibitory control on neuronal cell assemblies in the developing brain (Minlebaev et al., 2007; Kirmse et al., 2015) and counteract seizure generation (Woo et al., 2002; Nardou et al., 2011).

In the adult hippocampus new neurons are continuously generated throughout life. Similarly to the developing brain the GABA reversal potential is depolarized in newly generated hippocampal granule cells during the first 1-3 weeks after mitosis (Ge et al., 2006; Chancey et al., 2013; Heigele et al., 2016). Again, depolarizing GABAergic synapses do not generate hyperexcitability, but support sparse coding in the dentate gyrus (Lodge and Bischofberger, 2019). The newborn granule cells (GCs) improve learning and memory by enhancing the brains ability to discriminate similar memory items (Clelland et al., 2009; Creer et al., 2010; Sahay et al., 2011; Gu et al., 2012; Kheirbek et al., 2012; Bolz et al., 2015). Despite intense research, the underlying synaptic mechanisms are still largely unknown.

Newborn GCs integrate into the existing network via the formation of thousands of new glutamatergic and GABAergic synapses. The first glutamatergic synapses are formed as early as 1 week post mitosis (wpm) from hilar mossy cells followed by formation of afferent connections from the entorhinal cortex (EC) at 2 wpm (Deshpande et al., 2013; Chancey et al., 2014; Sah et al., 2017; Woods et al., 2018). Similarly, the first GABAergic interneurons were found to contact the young cells at around 1-2 wpm including soma-targeting parvalbumin (PV) positive basket cells as well as dendrite-targeting neurogliaform/ivy cells and somatostatin (SOM) interneurons (Ge et al., 2006; Markwardt et al., 2011; Alvarez et al., 2016; Groisman et al., 2020). Initially the GABAergic synapses show slow time course and small amplitude, followed by a maturational increase and speed-up of synaptic currents until about 6-8 wpm (Laplagne et al., 2006; Temprana et al., 2015; Groisman et al., 2020).

The smaller and slower GABAergic conductance in young cells was shown to facilitate synaptically evoked AP firing under certain conditions. Before 3 wpm the GABA_A_R reversal potential is depolarized close to the AP threshold (E_GABA_ ≈ -40 mV, Overstreet-Wadiche et al., 2005; Ge et al., 2006; Heigele et al., 2016). Therefore, low GABAergic activity can facilitate AP firing by coincidently active glutamatergic synapses, which would be subthreshold otherwise (Heigele et al., 2016). By contrast, larger GABAergic activity effectively blocks AP firing via shunting inhibition. At 4 weeks after mitosis, E_GABA_ is more hyperpolarized, but the GABAergic synapses are still not fully mature and may allow for more spiking in young cells (Marín-Burgin et al., 2012; Dieni et al., 2013). Remarkably, behavioral experiments show that young GCs are nevertheless only sparsely active (2-4%) during hippocampus-dependent learning (Kee et al., 2007; Danielson et al., 2016; Krzisch et al., 2016). The network mechanisms underlying this sparse activity are still largely unclear.

Although these studies indicate that GABAergic synapses can effectively inhibit AP firing in young GCs with depolarizing E_GABA_, it largely remains unclear how the different interneuron subtypes and their GABAergic synapses can achieve this. Using optogenetic stimulation of SOM- and PV-interneurons during whole-cell patch-clamp recordings together with computational modeling, we show that effective block of AP firing in young neurons is due to expression of synaptic α5-GABA_A_Rs. The α5-GABA_A_Rs-mediated currents showed a distinct nonlinear voltage-dependence, generating 4-times larger inhibitory conductance around the AP threshold as compared to the resting membrane potential. These unique properties of α5-GABA_A_Rs support the sparsification of AP firing in young GCs, even with depolarized GABA reversal potentials.

## Results

### PV-interneuron synapses activate outward rectifying GABA_A_ receptors in young granule cells

To examine the functional properties of PV- and SOM-interneuron mediated inhibition onto adult-born granule cells (GCs) of the dentate gyrus, we used optogenetic stimulation of different interneuron subtypes and performed whole-cell patch-clamp recordings in acute hippocampal brain slices from 5- to 10-week-old animals.

To identify newly generated GCs we used transgenic animals expressing DsRed under the control of the doublecortin promoter (Brown et al., 2003; Couillard-Despres et al., 2006). The input resistance was further used to estimate the cell age based on previous studies (Heigele et al., 2016; Li et al., 2017), to distinguish 2-3 weeks old (2-8 GΩ) from 3-4 weeks old cells (0.5-2 GΩ). To selectively activate PV-positive soma-targeting interneurons (basket cells, axo-axonic cells), we used virus-mediated expression of channelrhodopsin-2 (ChR2). A floxed-ChR2 virus was injected into DCX-DsRed animals crossed with a PV-Cre mouse line. As expected, viral expression of floxed-GFP for control was restricted to interneurons forming an axonal ramification within the granule cell layer (Fig. 1A,B). This allowed for both the selective expression of ChR2 in identified interneuron subtypes and the visualization of newborn GCs between 1-4 wpm.

**Fig. 1.**
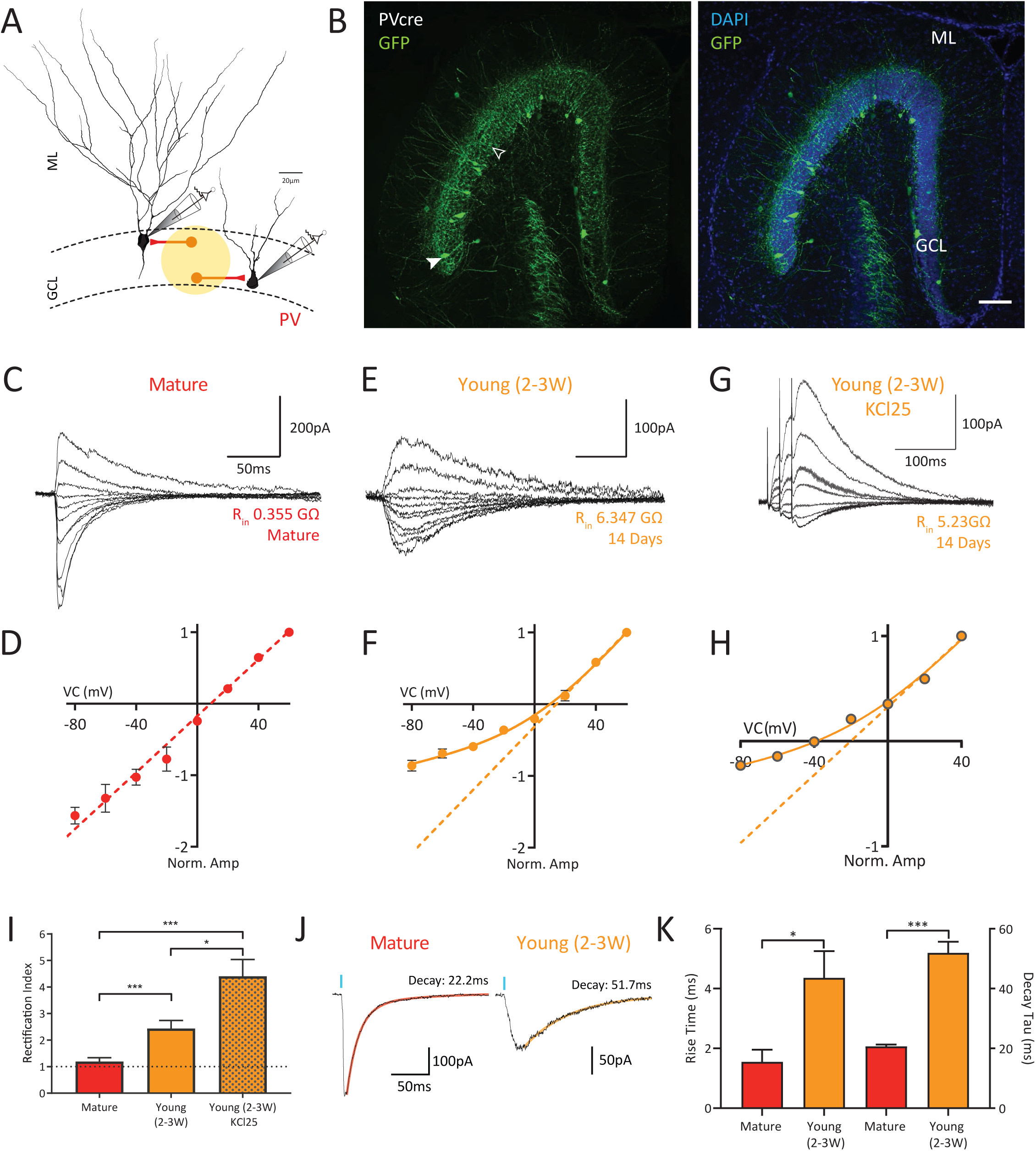
PV basket cell inputs onto young granule cells are non-linear and voltage dependent. **A**) Experimental design. PV-Cre x DCX-DsRed mice were injected with AAV-floxed-ChR2 in the ventral dentate gyrus (vDG). Optogenetic stimulation of the GCL was used excite PV-ChR2 -expressing basket cells forming synapses onto mature and young granule cell somata. Young 2- to 3-week-old granule cells were identified by the presence of DCX-DsRed fluorescence as well as their input resistance (2-8 GΩ). **B**) Confocal image of GFP-labelled basket cells after injection of AAV-floxed-GFP in the vDG of PV-Cre mice. Filled arrowhead: soma of a PV-basket cells; open arrowhead: basket cell axons in the granule cell layer (GCL). Scale bar 100µm. **C-D**) Light-evoked GPSCs recorded from mature GCs in symmetrical chloride conditions. Current amplitudes normalized to values at +60 mV. **E-F**) Light-evoked GPSCs recorded from DCX-expressing young GCs in symmetrical chloride conditions. Estimated age of the cell is indicated. Current amplitudes normalized to values at +60 mV. **G-H**) GPSCs evoked by electrical stimulation in the GCL were recorded from DCX-expressing young GCs in physiological chloride conditions (KCl25) in the presence of NBQX (10µM) and AP5 (25µM). A brief burst of 3 pulses (50Hz) was used to produce a larger inward current at -80mV. Current amplitudes normalized to values at +40 mV. Dashed lines in D, F and H represent linear fits to the outward currents. **I**) Rectification index in mature GCs (*n*=10) was significantly smaller than in young GCs (*n*=10) in symmetrical chloride, as well as in young GCs using physiological chloride concentration (*n*=8). **J-K**) Rise and decay time constants of light-evoked GPSCs were significantly slower in young versus mature GCs (*n*=8 and *n*=7). Significance indicated in figure as *: P<0.05, **: P< 0.01, *** P<0.001.

We recorded GABAergic postsynaptic currents (GPSCs) in response to ChR2-mediated light activation at different holding potentials in symmetrical chloride conditions (∼140 mM, Fig. 1CD). Similar to what was found in CA1 pyramidal cells (Schulz et al. 2018), PV-interneuron-evoked GABAergic inputs onto mature GCs showed a linear current-voltage relationship resembling classical GABA_A_R function. Surprisingly, PV inputs onto young GCs showed highly nonlinear GABAergic currents with much larger conductance at depolarized potentials relative to the resting membrane potential (Fig. 1E,F). To quantify the nonlinear amplification of conductance with depolarization, a rectification index (RI) was calculated as the ratio of the conductance obtained from the linear fit to the outward currents divided by the value measured at -80mV (Fig. 1I). The RI was significantly larger in young GCs at 2-3 wpm (RI=2.44±0.30, *n*=10) than in mature GCs (RI = 1.19±0.12, *n*=10, *P* ≤0.001). In young GCs at 3-4 wpm the rectification of PV-interneuron synapses was significantly smaller than in 2-3 wpm cells showing intermediate values (1.30±0.15, *n*=7, P=0.0097). Similarly, electrical stimulation of GABAergic fibers within the granule cells layer (GCL) in the presence of glutamate receptor blockers (10μM NBQX + 25μM AP5) revealed a linear response in mature GCs (RI = 1.14±0.08, *n*=8), while GABAergic synapses onto young GCs at 2-3 wpm were nonlinear (RI = 1.90±0.08, n=8, *P* ≤0.001, Supplemental Figure 1).

It is well known that young GCs show an elevated intracellular chloride concentration of 25 mM until about 3 wpm (Ge et al., 2006; Karten et al., 2006; Heigele et al., 2016). Therefore, we repeated the experiments above and measured soma-targeting GABAergic currents in young GCs using a physiological chloride concentration in the pipette solution. Under these conditions young cells showed even higher rectification, with a 4-fold larger outward conductance at depolarized potentials relative to resting-potential (RI=4.40±0.6, *n*=6, Fig. 1G,H). Similar to previous reports (Alvarez et al., 2016; Groisman et al., 2020), PV-IN inputs onto young GCs showed significantly slower rise time (young: 4.4± 0.9 ms, *n*=8 vs mature: 1.6±0.4 ms, *n*=7; *P* =0.029) and slower decay time course (young: 52.0± 3.7ms vs mature: 20.7± 0.6 ms; *P*=0.001) than synapses onto mature GCs (Fig. 1J,K). The different decay time course together with difference in voltage dependence indicates that young and mature GCs express different types of GABA_A_ receptors.

These results show that PV-interneuron-mediated inhibition in young and mature GCs is very different from each other. While soma-targeting GABA synapses in mature GCs show a classical linear behavior, GABAergic inhibition of young GCs is strongly dependent on the membrane potential, with a 4-fold amplification of conductance at depolarized potentials.

### SOM-interneuron inputs onto young and mature granule cells are outwardly rectifying

The nonlinear PV-interneuron-mediated synaptic currents in young GCs resemble properties of dendrite-targeting synapses from SOM-INs described in CA1 pyramidal cells (Schulz et al., 2018). To study GABAergic synapses formed by SOM-positive dendrite-targeting interneurons in the dentate gyrus (HIPP cells), we injected the floxed-ChR2 virus into DCX-DsRed animals crossed with a SOM-Cre mouse line (Fig. 2A,B).

Dendritic inputs onto young and mature GCs showed a similar nonlinear voltage-dependence in symmetrical chloride conditions (mature: RI = 2.59±0.47, *n*=8 vs young: RI=2.41±0.34, *n*=9, *P*=0.89; Fig. 2C-F). Similarly, using an internal chloride concentration of 25 mM revealed a high RI in SOM-IN evoked synapses in young GCs (RI=3.21±0.43, *n*=9, Fig. 2G,H). Interestingly, the rectification index from PV- and SOM-IN inputs onto young cells was very similar (Fig. 1I versus Fig 2I, KCL25: P=0.84), suggesting that the postsynaptic GABA_A_R composition in young GCs may be homogenous across the cell body and dendritic tree. In mature GCs however, the differences in rectification between different interneuron subtypes suggests that postsynaptic receptor composition is different in dendritic versus somatic synapses.

**Fig. 2.**
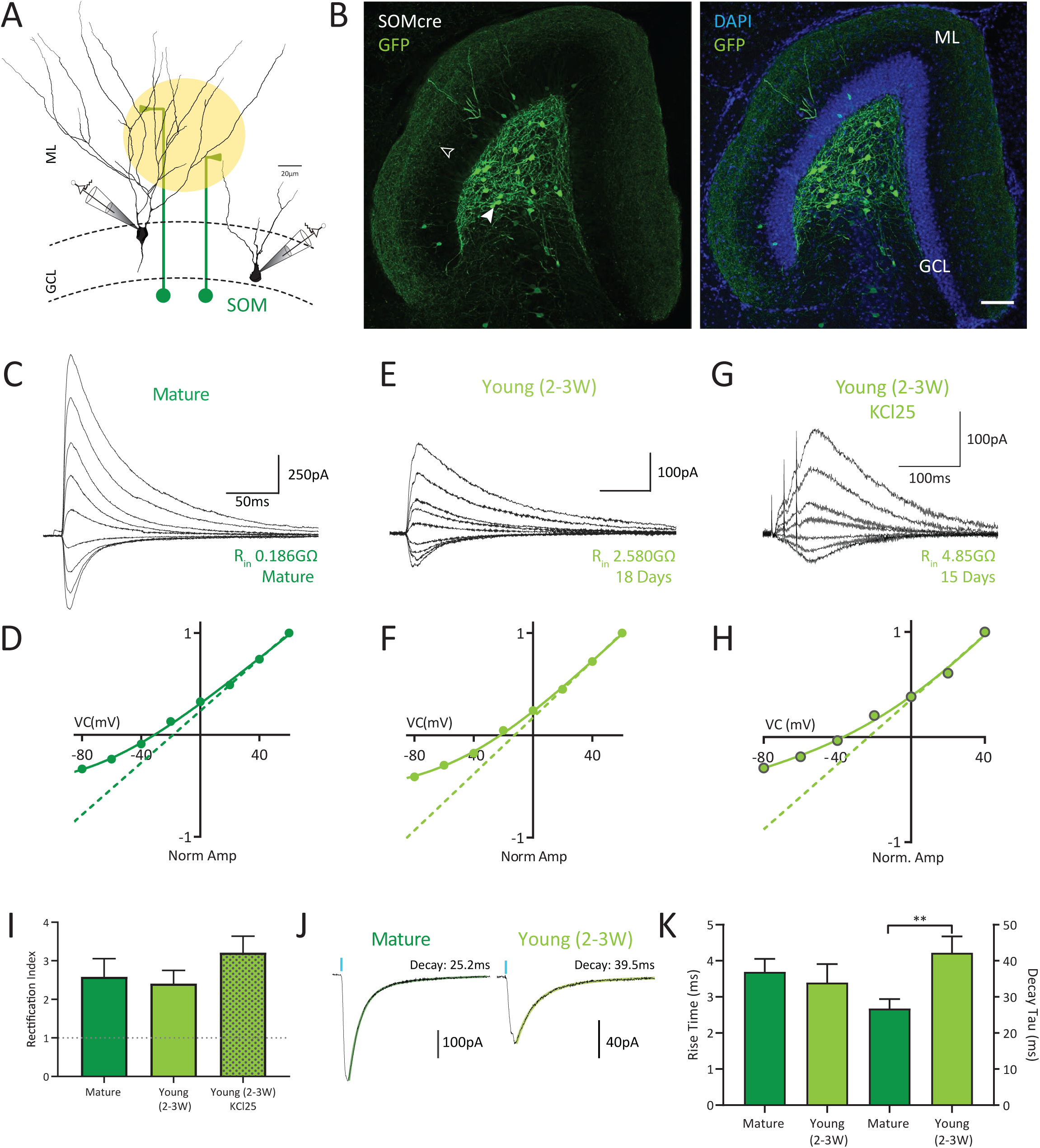
SOM HIPP-cell inputs are non-linear and voltage dependent in young as well as in mature granule cells. **A**) Experimental design. SOM-Cre x DCX-DsRed mice were injected with AAV-floxed-ChR2 in the ventral dentate gyrus (vDG). Optogenetic stimulation of the ML was used excite SOM-ChR2-expressing Hilar-perforant-path-associated (HIPP) interneurons forming synapses onto mature and young granule cell dendrites. Young 2- to 3-week-old granule cells were identified by the presence of DCX-DsRed fluorescence as well as their input resistance (2-8 GΩ). **B**) Confocal image of GFP-labelled HIPP cells after injection of AAV-floxed-GFP in the vDG of SOMcre animals. Filled arrowhead: soma of a SOM-expressing HIPP cells; open arrowhead: HIPP cell axons in the molecular layer (ML). Scale bar 100µm. **C-D**) Light-evoked GPSCs recorded from mature GCs in symmetrical chloride conditions. Current amplitudes normalized to values at +60 mV. **E-F**) Light-evoked GPSCs recorded from DCX-expressing young GCs in symmetrical chloride conditions. Current amplitudes normalized to values at +60 mV. **G-H**) GPSCs evoked by electrical stimulation in the ML were recorded from DCX-expressing young GCs in physiological chloride conditions (KCl25) in the presence of NBQX (10µM) and AP5 (25µM). A brief burst of 3 pulses (50Hz) was used to produce a larger inward current at -80mV. Current amplitudes normalized to values at +40 mV. Dashed lines in D, F and H represent linear fits to the outward currents. **I**) Similar rectification index in mature (*n*=8) and young GCs (*n*=9) in symmetrical chloride, as well as in young GCs in physiological chloride concentration (*n*=9). **J-K**) Rise and decay time constants of light-evoked GPSCs were significantly slower in young versus mature GCs (*n*=8 and *n*=9). Significance indicated in figure as *: P<0.05, **: P< 0.01, *** P<0.001.

Analysis of kinetic parameters revealed that dendritic inputs from SOM-INs did not show any difference in rise time between young and mature GCs (mature: 3.7± 0.4 ms, *n*=8; vs young: 3.4±0.5 ms, *n*=9, *P* =0.61, Fig. 2J,K). While the decay τ was slower in young than in mature neurons (young: 42.2±4.6 ms vs mature: 26.8±4.6 ms; *P*=0.006), the difference in kinetics was less pronounced than in perisomatic synapses (see Fig. 1K).

Taken together, the data show that young GCs express similar GABA_A_R in PV- and SOM-interneuron synapses with a remarkably nonlinear voltage dependence. During maturation, the soma-targeting synapses completely loose the rectifying voltage-dependence while SOM-interneurons continue to express nonlinear GABA_A_Rs.

### α5 subunit-containing GABA_A_Rs mediate synaptic inhibition onto adult-born granule cells

The different time course and voltage dependence of postsynaptic GABA_A_Rs in young versus mature GCs suggests that the postsynaptic GABA_A_R composition is different. α5 subunit-containing GABA_A_Rs have been shown to be present throughout the hippocampus including CA1 and dentate gyrus (Collinson et al., 2002; Serwanski et al., 2006; Zarnowska et al., 2009; Vargas-Caballero et al., 2010; Capogna and Pearce, 2011; Engin et al., 2015). Recently we have shown, that α5-GABA_A_Rs mediate synaptic inhibition onto apical dendrites of CA1 pyramidal cells and display strong outward rectification (Schulz et al. 2018). Therefore, α5-GABA_A_Rs were a good candidate for mediating the nonlinear GABAergic inhibition in GCs.

To test the contribution of α5-GABA_A_Rs we used RO4938581, a highly selective negative allosteric modulator (α5-NAM, 1μM; Ballard et al., 2009). Electrical stimulation in the granule cell layer (GCL) or outer molecular layer (OML) was used to target perisomatic or dendritic inputs, respectively, in the presence of NBQX and APV. While fast PSCs from GCL stimulation in mature GCs did not show a change in amplitude (α5-NAM: 100.6±4.6% of control, *n*=4, *P*=0.88; Fig. 3A), the amplitude of dendritic inputs was significantly decreased after application of the α5-NAM to 80.4±2.7% of control (*n*=9, *P*=0.004, Fig. 3BC). Furthermore, the decay τ for dendrite-targeting GABAergic inputs was decreased by α5-NAM application (69.4 ± 4.4ms to 57.6 ± 3.3ms, *P* =0.004), whereas soma-targeting inputs remained unchanged (Fig. 3D). This indicated that synaptic GABA_A_Rs in mature GCs show a similar profile to CA1 pyramidal cells.

**Fig. 3.**
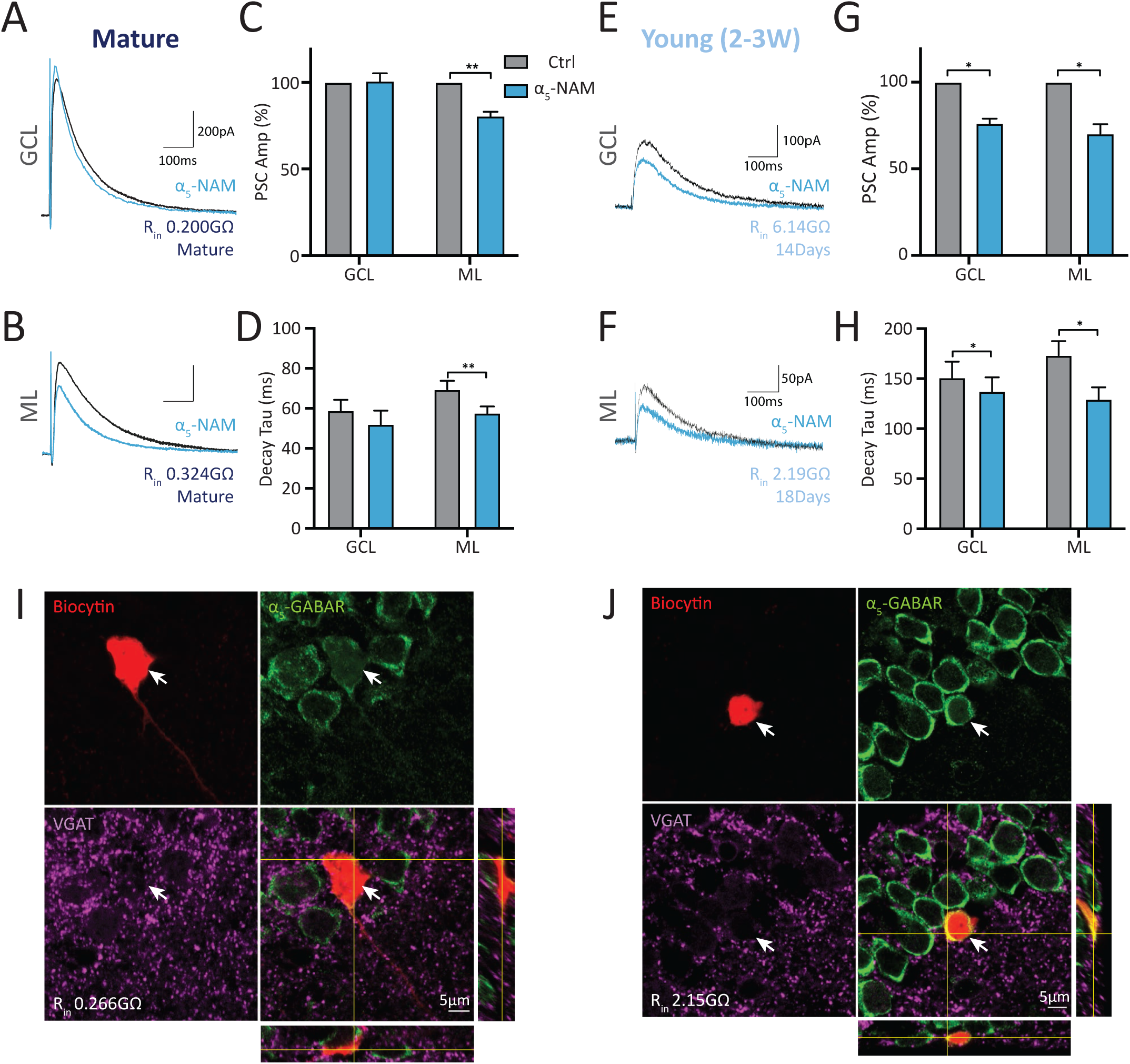
α5-GABA_A_Rs mediate dendritic and somatic Inhibition onto adult-born granule cells. **A-B**) Electrical stimulation of the GCL or ML was used to target somatic or dendritic inputs onto mature GCs, respectively. IPSCs were recorded at +60 mV in the presence of 10 µM NBQX and 25 µM APV using a CsCl-based pipette solution. **C**) IPSCs evoked by ML-stimulation were decreased in mature GCs after addition of the α5-NAM RO4938581 (1 µM, blue, *n*=9), in contrast to somatic IPSCs evoked by GCL stimulation (*n*=4). **D**) Similarly, α_5_-NAM reduced the decay time constant of dendritic IPSCs without changing somatic IPSCs. **E-F**) Representative mean IPSCs recorded from young GCs in a potassium-chloride-based pipette solution. Application of the α5-NAM (blue) shows that both dendritic and somatic inputs onto young GCs are mediated by α5-containing GABA_A_Rs. **G-H**) Both, GPSC amplitudes and decay time constants were significantly decreased in young GCs with ML-(*n*=6) as well as with GCL stimulation (*n*=6). **I**) Example of immunohistochemically labelled α5-GABA_A_Rs (green) and VGAT puncta (magenta) on a biocytin-filled (red) mature granule cell (266 MΩ). **J**) The same labelling as in I) on a 3-week-old young granule cell (2.15 GΩ).

In 2- to 3-week-old GCs, both, somatic and dendritic inputs were reduced after α5-NAM application. PSC amplitudes from GCL stimulation decreased to 76.3% ± 2.7% (*n*=6, *P*=0.031), while dendritic synaptic currents evoked by OML stimulation decreased to 70.2% ± 2.7% of control (*n*=6, *P*=0.031, Fig. 3E-G). Similarly, the somatic decay τ was reduced from 157.2 ± 16.0 ms to 137.4 ± 14.0 ms (*P*=0.016) and the dendritic decay τ from 173.4 ± 14.2 ms to 129.4 ± 12.0ms (*P* =0.031, Fig. 3H). RO4938581 has a maximal efficacy of only about 50% (Ballard et al., 2009). Thus, the contribution of α5-GABA_A_Rs might be much larger than the measured effect, suggesting that GPSCs in 2-3 week old GCs were largely generated by nonlinear GABA_A_ receptors containing α5-subunits. The comparable effect of α5-NAM on both perisomatic and dendritic inputs in young GCs indicated that GABA_A_R distribution may be homogeneous across the cellular compartments in striking contrast to mature GCs.

Biocytin filling and immunohistochemical staining of mature and young GCs was used to test for the presence of the α5-GABA_A_R combined with a marker for GABAergic presynaptic terminals (VGAT). Mature GCs did not have strong α5_-_labelling in the soma, consistent with the pharmacological data in Fig. 3A. However, VGAT puncta were present around the cell body and proximal dendrites (Fig. 3I). In contrast, young GCs had robust α5_-_GABA_A_R labelling throughout the soma, as well as surrounding VGAT-positive puncta (Fig. 3J). Taken together, these results suggest that GCs within the dentate gyrus express synaptic α5-GABA_A_Rs, with mature cells selectively expressing them in dendritic synapses, while young GCs show a homogenous distribution across both dendritic and somatic synapses.

### In mature GCs nonlinear GABA_A_ receptors are present in the extrasynaptic membrane of perisomatic synapses

Although it was shown that α5-GABA_A_Rs can contribute to phasic synaptic transmission at dendrite-targeting synapses (Serwanski et al., 2006; Ali and Thomson, 2008; Zarnowska et al., 2009; Schulz et al., 2018), they are traditionally considered to be extrasynaptic receptors (Farrant and Nusser, 2005). As rectification was not present in somatic synapses onto mature GCs, we sought to test whether GABA spillover from the synaptic cleft could potentially activate extrasynaptic α5-GABA_A_Rs. In order to increase GABA spillover, we inhibited the GAT1 transporter using NO711 (1μM), which normally clears GABA from the extrasynaptic space (Overstreet and Westbrook, 2003; Kersanté et al., 2013) and measured perisomatic inputs in young and mature GCs (Fig. 4). To prevent changes in GABA release, we additionally blocked presynaptic GABA_B_ receptors using CGP54626 (1μM).

**Fig. 4.**
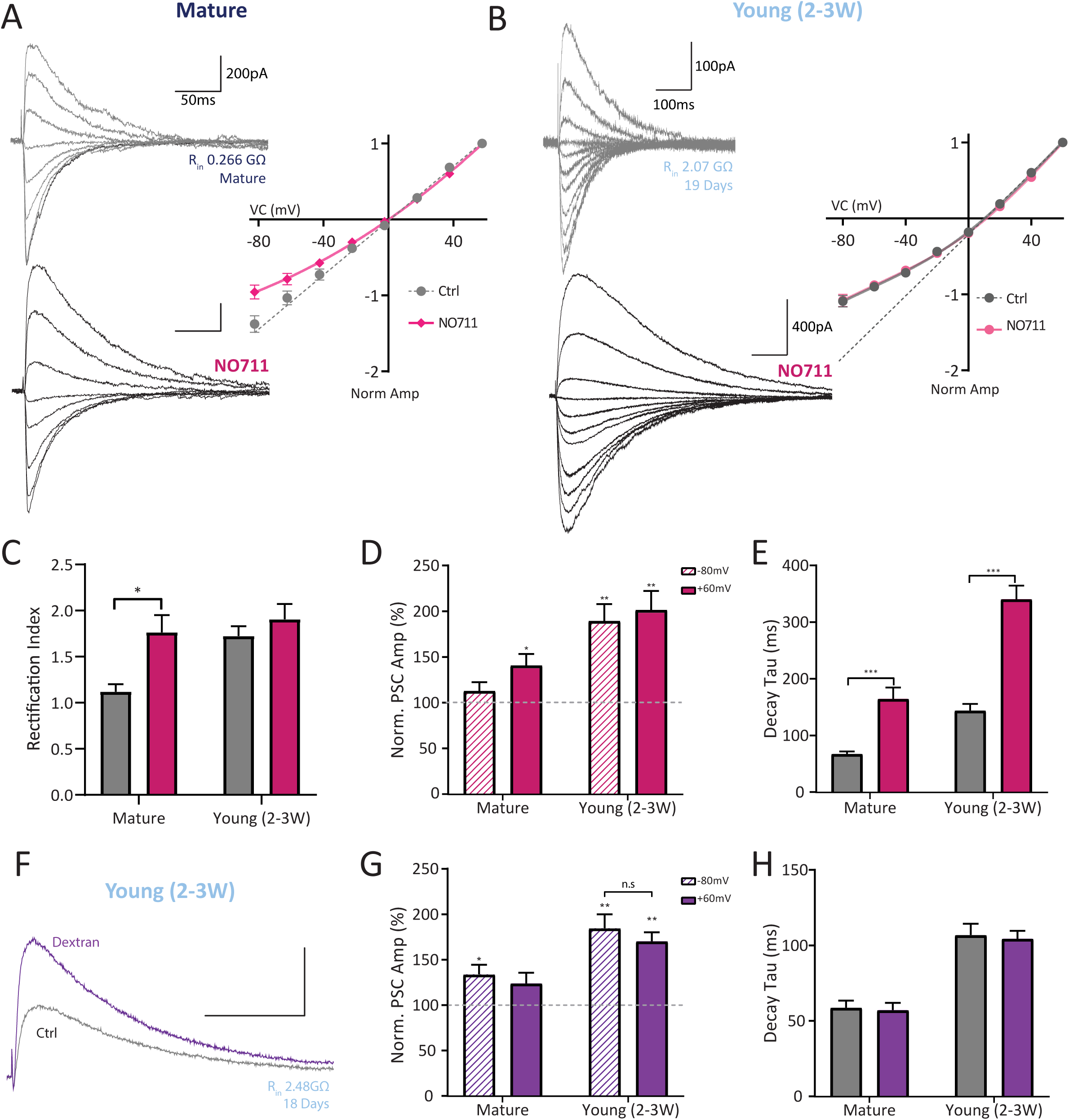
Increasing peri-synaptic GABA_A_R activation increases rectification in mature GCs. **A**) Electrical stimulation of the GCL was used to target somatic GABAergic inputs onto mature GCs (with 1μM CGP545626, 10μM NBQX, 25μM APV). The GABA reuptake was blocked using NO711 (1μM) to increase peri- and extra-synaptic receptor activation (top, control in grey, bottom NO711 in black). IPSCs were recorded at increasing membrane potentials (−80mV to +60mV) in symmetrical chloride conditions. Outward GPSCs in mature cells showed larger increase in amplitude than inward currents. As a consequence, the current-voltage relationship becomes nonlinear post NO711 (inset, red, *n*=13). **B**) GPSCs from young GCs showed a large increase in amplitude of both inward and outward currents, producing similar rectification post-NO711 application (*n*=11). **C**) The rectification index of mature GCs significantly increases with more spillover and extrasynaptic receptor activation (*n*=13), while in young GCs this remains the same (*n*=11). **D**) The outward PSCs increased significantly with NO711 application (filled bar), whereas the synaptic inward currents (dashed bar, left) remained largely unchanged in mature GCs. In yGCs, both outward and inward currents were significantly increased. **E**) The decay τ of both mature and young GCs were significantly increased. **F**) 5% dextran was used to confirm synaptic transmission in yGCs. Example traces of IPSCs recorded at +60mV show a large increase in amplitude and similar decay τ with dextran wash-in. **G**) Young and mature GCs showed a significant increase in the normalized amplitude with dextran application (*n*=9 and *n*=9). **H**) Decay time constants, however, were not significantly slowed by Dextran, suggesting that the currents in young and mature cells are mediated via postsynaptic GABA_A_ receptors.

Amplitudes and the decay τ of recorded PSCs were increased in both mature and young GCs, consistent with the block of GABA reuptake (Fig. 4A-E). Interestingly, mature GCs showed a larger increase in GABAergic outward currents as compared to inward currents (139.3±13.9%, *P*=0.033 vs 111.2±11.3%, *n*=13, *P*=0.41). As a consequence the current-voltage relationship changed from linear towards outward-rectifying (Fig. 4A, magenta NO711 vs grey control) and the rectification index was significantly increased from 1.12± 0.08 to 1.76± 0.18 (*n*=8, *P* =0.016; Fig. 4C). This indicated that spillover in mature GCs indeed activates nonlinear extrasynaptic GABA_A_Rs producing outward rectification, similar to what has been reported by Pavlov et al. (2009) for extrasynaptic GABA_A_Rs in CA1 pyramidal neurons.

In 2- to 3-weeks-old GCs, inhibition of GAT1 transporters resulted in a uniform increase in amplitude (188.0± 19.85% inward, *n*=11, *P*=0.002 vs 199.9± 22.29% outward, *P*=0.002) across all holding potentials (Fig. 4B), which did not change the voltage dependence (RI=1.73± 0.10 in control vs. RI=1.90± 0.16 in NO711, *P*=0.25; Fig. 4C). The stability of the rectification index in young cells with increased spillover indicated that young GCs may have homogeneous receptor distribution between the synaptic and extrasynaptic cell membrane.

As rectification in young GCs was not changed with NO711, we sought to confirm the presence of true synaptic transmission and to ensure that GABAergic signaling onto young GCs is not only mediated by spillover from neighboring GABAergic synapses onto mature GCs. To slow down diffusion of GABA out of the synaptic cleft we used dextran, a branched molecule with high molecular weight, which increases the viscosity of the extracellular space (Nielsen et al., 2004; Markwardt et al., 2009). This effectively reduces spillover from the synapse while enhancing the synaptic concentration of neurotransmitter. The decreased mobility of GABA and thus higher concentration within the synaptic cleft increases the occupation and activation of both pre- and postsynaptic receptors. In order to prevent changes in GABA release we blocked presynaptic GABA_B_ receptors using CGP54626 (1μM). If newborn GCs only received GABAergic inputs from spill-over of neighboring mature synapses we would expect to see a decrease in GPSC amplitude with dextran application. In contrast, if young GCs receive synaptic inputs, the amplitudes will remain the same or increase with dextran.

Application of dextran clearly increased the amplitude of synaptic currents in young as well as in mature GCs (Fig. 4F,G). While the amplitude of inward currents increased to 133.8±10.7% (*n*=9, *P*=0.014) in mature GCs, the increase in young GCs was even larger (184.6±15.3%, *n*= 9, *P*=0.004, Fig. 4G). This is consistent with a higher concentration of GABA within the synaptic cleft due to reduced diffusion. This increase in GPSC amplitude by dextran directly supports the notion that young GCs indeed receive true synaptic inputs, and not just spillover from neighboring synapses. Furthermore, the large increase in amplitude in 2- to 3-weeks-old GCs suggests that postsynaptic GABA receptors are not saturated. This suggests that application of dextran increased receptor occupancy via an increased GABA concentration within the synaptic cleft.

Kinetic analysis of GABAergic currents showed that the rise time of GPSCs were not changed after dextran application (+60mv, mature GCs 2.1±0.5ms to 2.0±0.5ms, *n*=9 *P*=0.84; young GCs 6.0±0.5 to 5.5±0.3, *n*=9, *P*=0.20). Furthermore, there was no effect on decay τ in young and mature GCs (Fig. 4H). In case GPSCs had been mediated by extrasynaptic GABA_A_Rs, we would have expected an increase in rise time with dextran application, which was not observed. Finally, as dextran application increased inward and outward currents in young GCs equally (inward to 184.6 ± 15.3%, outward to 170.2 ± 10.1%, *n*=9, *P*=0.30, Fig. 4G), there was no change in the magnitude of outward rectification. Thus, voltage-dependent nonlinear GABA_A_Rs are present in synapses of young GCs, but are excluded from the postsynaptic membrane of perisomatic synapses in mature GCs.

### α5-GABA_A_Rs in young GCs promote NMDAR-mediated excitation

What is the functional role of synaptic α5-GABA_A_Rs in adult-born GCs? As synaptic α5-GABA_A_Rs have slow kinetics similar to that of NMDARs (Schulz et al., 2018, 2019), we wanted to know if the slow GABAergic depolarisation mediated by these receptors helps to facilitate NMDAR activation in young GCs. To test whether α5-GABA_A_Rs modulate NMDAR activation, we performed whole-cell current-clamp recordings in the absence of any blockers. To maintain the physiological chloride reversal potential of -35 mV in young GCs (<3 wpm) we used a potassium gluconate-based pipette solution with 25 mM chloride (Heigele et al., 2016; Li et al., 2017).

By stimulating in the molecular layer (brief burst of 3 pulses at 50 Hz) we were able to effectively recruit glutamatergic and GABAergic inputs onto young GCs resulting in subthreshold PSPs with an average depolarization of 35.3±1.0 mV (*n*=12; Fig. 5A). Bath application of the α5-NAM (1μM) resulted in a large decrease of the burst PSP reducing the integral to 56.2±8.8% of control (*n*=7, *P*=0.008; Fig. 5A,B). Subsequent wash-in of AP5 (50μM) resulted in a further small NMDA-dependent decrease by about 15% relative to the baseline integral (40.8±5.9% of baseline, *P*=0.008). By contrast, application of AP5 in ACSF caused a 32% reduction of the integral to 68.0±2.9% of control (*n*=9, *P*=0.003; Fig. 5C,D). The 2-fold larger NMDA component in ACSF (32.0± 2.9%, *n*=9) versus in α5-NAM (14.4±4.4%, *n*=8, *P*=0.008; Fig. 5E) indicated that GABAergic depolarization via α5-GABA_A_Rs facilitates NMDA receptor activation. This is easily explainable by the voltage-dependent magnesium block of the NMDARs.

**Fig. 5.**
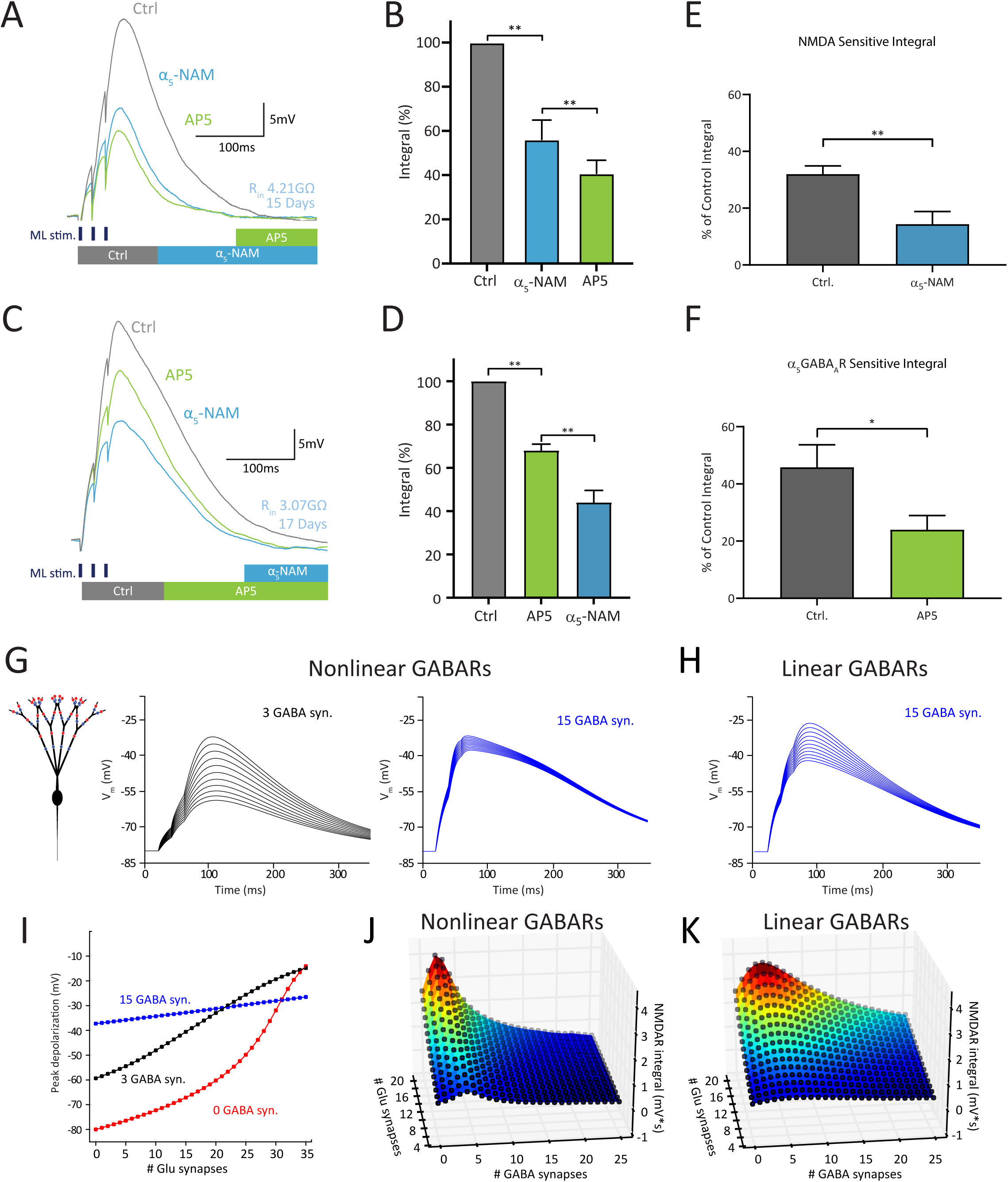
Activation of depolarizing α5-GABA_A_Rs controls NMDAR recruitment. **A**) A short burst (3@50Hz) of electrical stimulation of the GCL was used to evoke a PSP with an amplitude of about 35mV. The application of α5-NAM (1 µM), known to reduce specifically the conductance of α5-GABA_A_Rs by approx. 50%, strongly decreased the burst PSP (blue). The PSP after additional application of the NMDAR antagonist AP5 (50 µM) is shown in green. **B**) The PSP integral dramatically dropped with α5-NAM application, while further AP5 application produced a significant but modest reduction (*n*=7). **C-D**) AP5 application in control ACSF produced a large change in PSP integral, while subsequent application of α5-NAM resulted in a smaller decrease of the integral (*n*=9). **E**) The NMDA-sensitive integral was smaller in the presence of α5-NAM (*n*=9) than in control (*n*=8). **F**) Similarly, the α5-NAM-sensitive integral was larger in control than in the presence of APV, suggesting a nonlinear interaction between α5-GABA_A_Rs and NMDARs. G-K) Computational modelling. **G**) Distribution of GABAergic (blue) and glutamatergic synapses (red) on dendrites of a model cell approximating a newborn 2.5-week old GC (left panel). Brief-burst activation (3@50Hz) of an increasing number of glutamatergic synapses (1 to 21 in steps of 2) coincident with 3 GABAergic synapses with properties resembling the nonlinear α5-GABA_A_Rs as detailed in methods (middle panel). The same glutamatergic synapses coincident with 15 GABAergic synapses (right panel). **H**) The same stimulation pattern as in the right panel of (G) with linearized GABA_A_Rs resulted in a weaker shunt. **I**) The contribution of glutamatergic synapses to the burst PSP amplitude deceases with increasing number of nonlinear GABAergic synapses. **J**) The contribution of NMDARs to the burst-PSP integral is maximal with coincident small number of GABAergic synapses but is largely blocked with brief burst activity in more than 10 GABAergic synapses. **K**) The window with significant contribution of NMDARs to the burst-PSP integral widens if GABA synapses are linear.

Additional wash in of the α5-NAM in the presence of AP5 further reduced the PSP integral by 24% relative to baseline (to 44.04±5.5%, *P*=0.002; Fig. 5C,D). Comparing the effect of the α5-NAM after AP5 with its effect in ACSF (24% vs 44%, Fig. 5F), suggested that also the α5-GABA_A_Rs contribute to depolarization in a voltage-dependent manner. Therefore, NMDARs and α5-GABA_A_Rs interact in a highly nonlinear manner due to the voltage-dependent conductance of both receptor subtypes.

To mechanistically understand the voltage-dependent interaction of NMDAR and α5-GABA_A_Rs, we developed a compartmental cable model of a 2.5-week-old granule cell (Rin = 4GΩ, see methods). The model included GABAergic and glutamatergic synapses homogenously distributed along the dendritic tree and a depolarizing E_GABA_ (−35mV, Fig. 5G). Simulating the activation of an increasing number of glutamatergic EPSPs (3@50Hz) coincident with 3 depolarizing GABAergic SOM-like synapses generated large burst-EPSPs (Fig. 5G, left). In agreement with results from our physiological experiments, these burst EPSPs were larger than in absence of GABAergic synapses (Fig. 5I, 3 GABA vs 0 GABA syn.). However, increasing the number of GABAergic synapses to 15 shunted the effect of glutamate-dependent depolarization (Fig. 5I). This GABAergic shunting was due to the voltage-dependent increase of the α5-GABA_A_R-conductance, because the shunting effect was much smaller when SOM synapses were modelled with conventional linear GABA_A_Rs (Fig. 5H vs 5G, right). When we systematically mapped the effect of varying numbers of active GABA and glutamate synapses, the simulation results showed that the nonlinear gating of α5-GABARs promotes NMDAR activation within a restricted band of GABAergic activity (Fig. 5J). In contrast, linear GABA_A_Rs support NMDAR-activation over a much broader range (Fig. 5K). Therefore, the results from computational modelling indicated that depolarizing GABAergic synapses in young GCs can amplify NMDAR activation at glutamatergic synapses. However, the distinct functional properties of synaptic α5-GABA_A_Rs in young GCs restrict this GABAergic amplification to a narrow activity window.

Taken together, the data show that α5-GABA_A_Rs promote voltage-dependent GABAergic depolarization of young GCs, which in turn facilitate the activation of voltage-sensitive NMDARs. At the same time, large GABAergic activity can limit NMDAR-mediated depolarization by reducing the peak potentials via shunting inhibition.

### α5-GABA_A_Rs bidirectionally mediate excitation and shunting inhibition of adult born granule cells

Previously we have shown that GABAergic synaptic activity can block action potential (AP) generation in adult-born young GCs at 1-3 wpm via depolarizing shunting inhibition (Heigele et al., 2016). To investigate the contribution of α5-GABA_A_Rs to shunting inhibition, we performed whole-cell current-clamp recordings in the presence of NBQX and AP5 to block glutamatergic inputs (10 μM and 25 μM, respectively). Perisomatic GABAergic inputs were activated via local electrical stimulation of the GCL, using an intensity sufficient to evoke shunting inhibition (30-60 μA). Coincident mock EPSPs (approx. 20 mV) were generated via current injection to simulate postsynaptic integration of excitatory and inhibitory inputs (Fig. 6A).

**Fig. 6.**
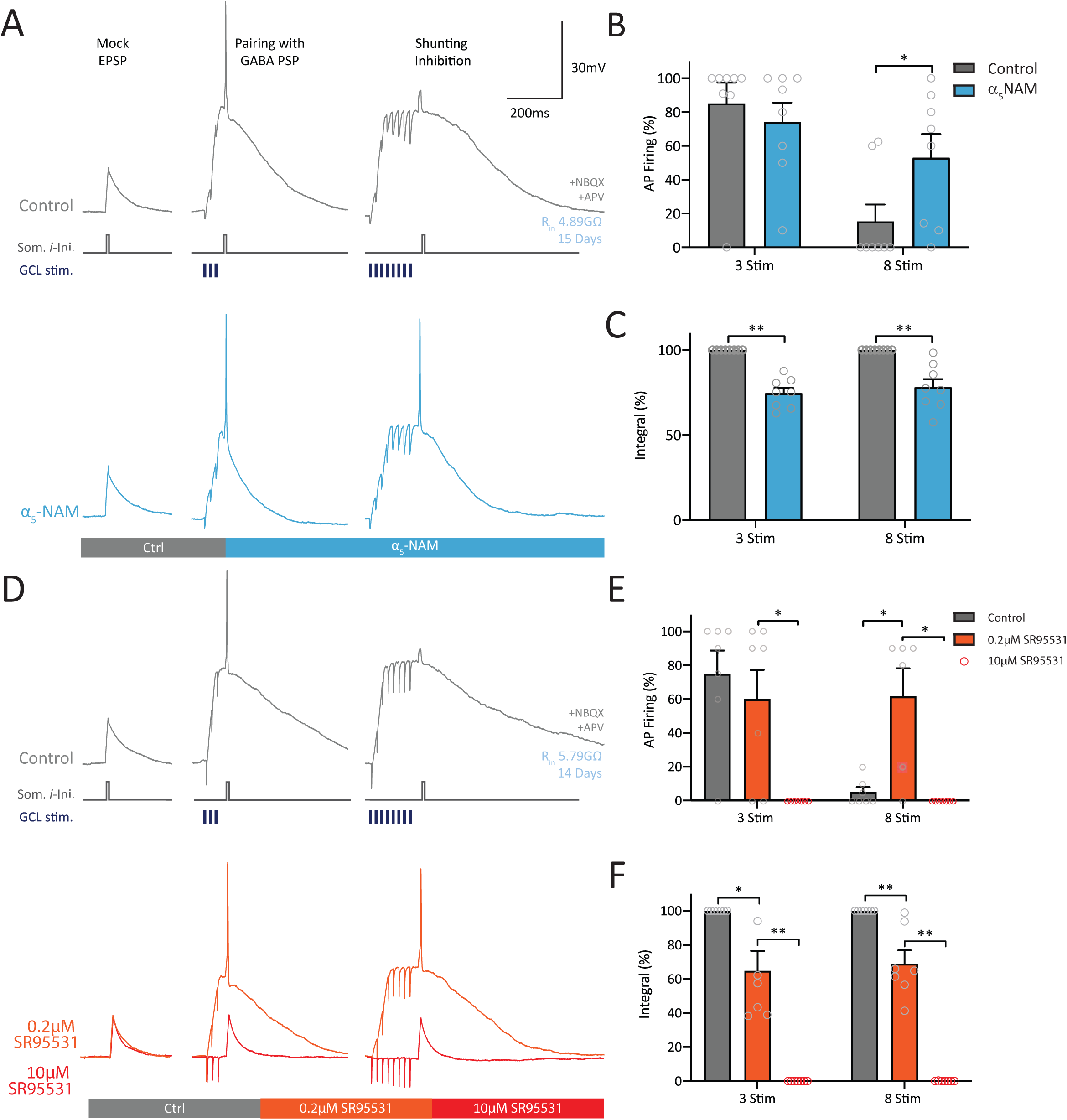
α5-GABA_A_Rs maintain sparse-firing in yGCs with shunting inhibition. **A**) Experimental design. A somatic current injection (10ms) was delivered to generate a mock PSP of about 20 mV, which was paired with a burst stimulation of 3 or 8 pulses in the GCL (3@50Hz or 8@50Hz) in the presence of NBQX (10μM) and APV (25μM). The stimulation intensity was adjusted to reliably produce shunting inhibition with the 8@50Hz burst stimulation (30-60µA). Example traces are shown in control (grey) and after the application of α5-NAM (1µM, blue). **B**) Mean AP firing probability of the 3 and 8@50Hz burst stimulation. AP firing with 3@50Hz was not significantly affected with α_5_-NAM application, while shunting inhibition with 8@50Hz was significantly reduced, resulting in a large increase in AP firing. Grey open circles represent individual cells. **C**) Mean PSP integrals show similar reduction with α5-NAM application. **D**) Example traces of control (grey) with sequential application of 0.2µM (orange) and 10µM (red) gabazine (SR95531). **E**) Similarly, to α5-NAM, 0.2µM gabazine did not affect AP firing with the 3-pulse burst, but significantly reduced shunting inhibition, resulting in a large increase of AP firing with the 8@50Hz burst. Subsequent application of 10µM gabazine (red open circles) abolished all AP firing. **F**) Mean integrals were reduced with 0.2µM gabazine, while 10µM gabazine fully blocked the remaining GPSPs.

Brief burst stimulation of perisomatic GABAergic synapses (3@50Hz) facilitated AP firing, while longer burst stimulation using the same stimulation intensity (8@50Hz) inhibited spiking via shunting inhibition similar to our previous report (Heigele et al., 2016). The application of the α5-NAM (1μM) had a differential effect, leaving AP firing probability in response to the short burst largely unaffected (85.1±12.3% in control vs 74.2±11.4% in α5-NAM, *n*=8, *P*=0.15). By contrast, shunting inhibition was strongly reduced with a significant increase in AP firing probability from 15.3±10.0% to 53.0±13.9% during the long burst (*n*=8, *P*=0.016; Fig. 6B). Both, the 3- and 8-pulse stimulation responses showed a decrease of the PSP integral (3 stim 74.6± 3.1%, *P* =0.004; 8 stim 78.1± 4.7%, *P* =0.004) as expected after a reduction in GABAergic depolarization. Similar results were obtained with the stimulation electrode located in the molecular layer to activate dendrite-targeting interneurons (data not shown). These observations suggested that α5-containing GABA_A_Rs effectively mediated shunting inhibition, while having only a relatively mild effect on GABAergic excitation. This is consistent with the biophysical properties of the α5-containing GABA_A_Rs in young GCs showing an about 4-fold larger conductance around the AP threshold (−35mV) compared to -80mV (see Fig. 1). Thus, blocking α5_-_GABA_A_Rs widens the window of opportunity for AP discharge in young cells and increases overall excitability.

Previously we reported that a low concentration of gabazine (SR95531, 0.2μM) unspecifically blocks about 50% of GABA_A_Rs in newly generated young GCs (Heigele et al., 2016). Thus, we sought to test whether a low concentration of gabazine had a similar effect to that of α5-NAM. In agreement with this prediction, we found that low gabazine (0.2μM) had little effect on GABAergic excitation as shown by the similar AP probabilities with short bursts (from 79.5± 10.9% to 61.2± 13.7%, *n*=9, *P*=0.38, Fig. 6D,E). Furthermore, shunting inhibition was strongly reduced, allowing effective AP firing from 4.38± 2.6% in control to 69.6± 13.3% in gabazine (*n*=8, *P*=0.031). Integrals for both the 3-stim and 8-stim bursts were significantly decreased (3 pulses: 63.1±9.1%, *P*=0.012; 8 pulses: 68.9±6.1%, *P*=0.004, Fig. 6F) similar to α5-NAM. Subsequent application of 10μM gabazine was sufficient to fully block any remaining PSP, showing that all synaptic responses were indeed GABA_A_R-mediated potentials (Fig. 6D-F). This suggests that newly generated young GCs express a functionally homogenous population of synaptic GABA_A_Rs along the somato-dendritic domain, consistent with a dominant contribution of α5-GABA_A_Rs.

Together, these results show that a 25-50% reduction of synaptic GABA_A_R-conductance in young GCs is sufficient to relieve shunting inhibition and restore AP firing with strong stimuli. Therefore, the 4-fold voltage-dependent increase in GABA conductance appears to be crucial to provide effective shunting inhibition at depolarized potentials close to the AP threshold (−35 mV).

### Voltage-dependent inhibition promotes sparse coding in young adult-born granule cells

To test the hypothesis that voltage-dependent amplification of the α5_-_GABA_A_R-conductance controls synaptic recruitment of young GCs, we again utilized computational modeling. This allows to systematically vary the number and change the functional properties of the activated synapses, which cannot be achieved by currently available experimental techniques. We simulated AP firing in young GCs in response to a brief burst stimulation (3@50Hz) of a varying number of EC fibers and SOM interneuron synapses (Fig. 7). Using realistic properties of glutamatergic and GABAergic synapses in 2.5 wpm GCs (see methods) a brief burst of up to 20 glutamatergic synapses did not generate APs (Fig. 7A, left). By contrast, pairing subthreshold glutamatergic EPSPs with 10 active SOM synapses reliably elicited an AP, while spiking was largely blocked with 2-times more GABA synapses (Fig. 7B). This nicely captured the experimental observation that depolarizing GABA synapses can facilitate AP firing in young GCs. However, similar to the mock EPSP above (Fig. 6), the synapse-evoked generation of APs was restricted to a narrow activity window (Fig. 7C). By contrast, when nonlinear α5-GABA_A_Rs in SOM synapses were replaced by conventional linear GABA_A_Rs, GABAergic excitation of young GCs was much more broadly tuned (Fig. 7D,E).

**Fig. 7.**
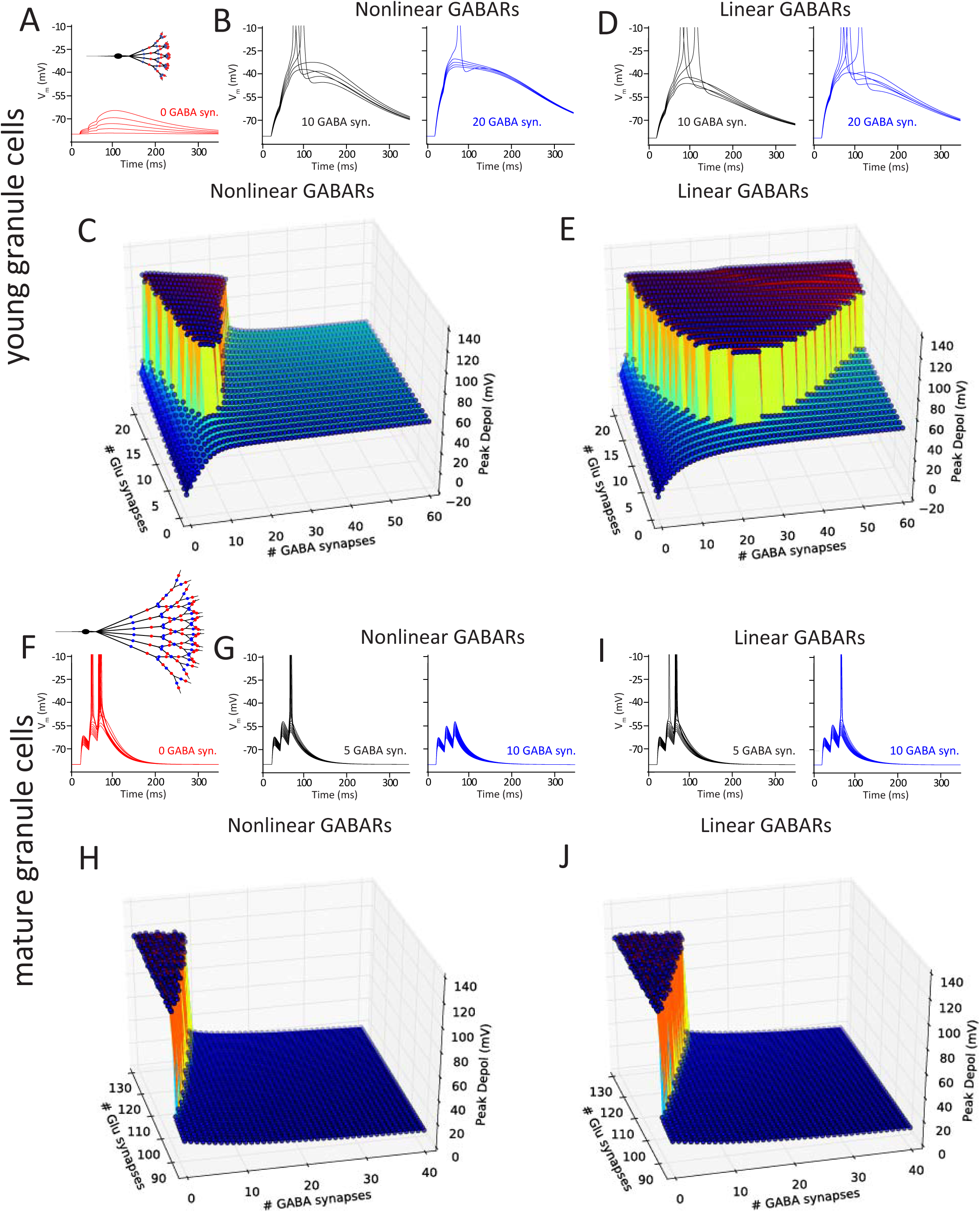
Voltage-dependent inhibition promotes sparse coding in young adult-born granule cells. **A**) Simulation of burst PSPs (3@50Hz) in a young GC with 1, 5, 9, 13 and 17 active glutamatergic synapses. GABAergic synapses are inactive. Inset: Distribution of GABAergic (blue) and glutamatergic synapses (red) on dendrites of the newborn 2.5-week-old model cell. **B**) Coincident activation of 10 (black) but not 20 GABAergic synapses (blue) facilitate AP firing in the young GC. **C**) Simulation of burst-induced AP firing with different combinations of active glutamatergic and GABAergic synapses, indicate that spiking is restricted to low interneuron activity. More than 15-20 active GABAergic synapses effectively block spiking in young GCs. **D-E**) When the nonlinear outward rectification of GABAergic synapses is replaced by linear GABA_A_Rs, spiking occurs much more widespread (E). **F**) Simulation of burst PSPs (3@50Hz) in a mature GC with 85 to 129 (in steps of 4) active glutamatergic synapses. GABAergic synapses are inactive. Inset: Distribution of GABAergic (blue) and glutamatergic synapses (red) on dendrites of the model cell. **G**) Coincident activation of 5 (black) and 10 GABAergic synapses (blue) reduce AP firing in mature GCs. **H**) Simulation of burst induced AP firing with different combinations of active glutamatergic and GABAergic synapses, indicate that spiking is restricted to low interneuron activity. More than 10 active GABAergic synapses effectively block spiking in mature GCs. **I-J**) In case the nonlinear outward rectification of dendritic GABAergic synapses is replaced by linear GABA_A_Rs, inhibition of spiking is less effective (J).

As mature GCs also express α5_-_GABA_A_Rs in SOM synapses, we wondered whether the nonlinear voltage-dependent inhibition has a similar effect on AP output in mature neurons. Therefore, we used a compartmental model of a mature GC based on previously established parameters (Schmidt-Hieber et al., 2007; Schmidt-Hieber and Bischofberger, 2010). In contrast to the young GCs, activation of GABAergic synapses always reduced AP output due to the negative E_GABA_ (−75mV, Fig. 7F,G). Therefore, EPSP-evoked spiking was effectively controlled by an increasing number of active SOM interneurons, which mimics lateral feedback inhibition. When α5-GABA_A_Rs were replaced with linear GABA_A_Rs, firing was more broadly tuned (Fig. 7J vs 7H), although this effect was a lot less dramatic than in young GCs. This shows that, although the voltage-dependent increase of the α5-GABA_A_R-conductance controls dendritic NMDAR activation and spiking in mature neurons (Schulz et al., 2018), linear inhibition is also effective. In young GCs, however, when E_GABA_ is depolarizing, GABAergic outward rectification is crucial for any inhibition of AP firing.

Together, these results show that properties of postsynaptic GABA_A_Rs in adult-born young GCs facilitate NMDAR activation and AP firing during small amplitude GABAergic PSPs coincident with glutamatergic EPSPs. However, due to their 4-fold voltage-dependent increase in conductance, the α5-GABA_A_R can powerfully shunt AP initiation. This mechanism allows for effective GABAergic inhibition of AP firing with enhanced GABAergic interneuron activity, supporting sparse AP firing in mature as well as in newly generated young GCs.

### Discussion

The analysis of SOM- and PV-interneuron-mediated synaptic inhibition onto adult-born GCs, provided several important insights. First, parvalbumin-positive interneurons activate synaptic α5-GABA_A_Rs in the perisomatic region of young GCs at 2-3 weeks post mitosis, while these receptors are excluded from perisomatic synapses in mature GCs. The α5-GABA_A_Rs in young GCs show distinct outward rectification with a 4-fold larger conductance at the AP threshold relative to the resting membrane potential. Second, somatostatin-interneurons also activate α5-GABA_A_Rs already at 2-3 wpm and continue to activate these receptors in dendritic synapses of GCs throughout maturation. Third, due to the depolarizing E_GABA_ in young neurons, activation of a low number of synaptic α5-GABA_A_Rs facilitate the opening of NMDARs during coincident activation of glutamatergic synapses. In spite of the depolarizing E_GABA_, strong activation of synaptic α5-GABA_A_Rs can nevertheless inhibit AP firing in young neurons, similar to mature GCs, because of the voltage-dependent conductance profile.

Nonlinear α5-GABA_A_Rs may not only be important for newborn granule cells in the adult brain, but also control firing of immature neurons during forebrain development. The α5-subunit of GABA_A_Rs is strongly expressed in the developing neocortex and hippocampus with a peak at around P3-P7 (Laurie et al., 1992; Poulter et al., 1992; Liu et al., 1998; Brooks-Kayal et al., 2001). While the expression of α5-GABA_A_Rs remains high in the hippocampus, the expression in the neocortex subsequently decreases towards lower adult levels. In mouse somatosensory cortex the expression of α5-GABA_A_Rs correlates with the slow decay of spontaneous GABAergic synaptic currents at P7 (Dunning et al., 1999). Remarkably, early network activity in P1-P7 mouse cortex is constrained by inhibitory GABAergic synapses, with a major contribution of α5-GABA_A_Rs (Minlebaev et al., 2007; Kirmse et al., 2015; Valeeva et al., 2016). Furthermore, GABAergic synapses can protect the early postnatal brain from epileptic seizures instead of inducing them (Woo et al., 2002; Nardou et al., 2011). The voltage dependence of these early postnatal α5-GABA_A_Rs has not been investigated. However, our results suggest that the inhibitory effect of postnatal GABAergic synapses is at least partially mediated by outward rectification of α5-GABA_A_Rs, which promote effective shunting inhibition in immature neurons with depolarized E_GABA_ and prevent seizures.

Although α5-GABA_A_Rs were previously considered to be mainly located on the extrasynaptic membrane, there is mounting evidence for synaptic localization (Serwanski et al., 2006; Ali and Thomson, 2008; Zarnowska et al., 2009; Schulz et al., 2018, 2019; Magnin et al., 2019). While α5-GABA_A_Rs can bind to extrasynaptic radixin they can also bind to postsynaptic gephrin and can thereby shuttle between synaptic and extrasynaptic sites in a phosphorylation-dependent manner (Brady and Jacob, 2015; Hausrat et al., 2015; Jacob, 2019). As our dextran data indicate that extrasynaptic and synaptic GABA_A_Rs in young GCs share the same properties (Figure 4), the available α5-GABA_A_Rs might dynamically shuttle between synaptic and extrasynaptic sites. For developing hippocampal neurons *in vitro* it was reported that the synaptic pool of α5-GABA_A_Rs is specifically important for spine maturation (Brady and Jacob, 2015). During later neuronal maturation, however, α5-GABA_A_Rs appear to be progressively excluded from perisomatic synapses in GCs as well as in pyramidal cells (Zarnowska et al., 2009; Schulz et al., 2018), probably due to expression of GABA_A_ receptors with higher affinity for gephrin (Brady and Jacob, 2015). As cell-autonomous deletion of α5-GABA_A_Rs in newly generated young GCs has been shown to affect dendritic growth (Deprez et al., 2016), a more detailed analysis of the developmental role of synaptic versus extrasynaptic GABA receptors in newborn GCs of the adult hippocampus is an important topic for future studies.

According to our experimental and computational modelling results, the synaptic α5-GABA_A_Rs appear to have a large impact on synaptic recruitment and AP firing in mature as well as in newborn GCs from 2 wpm onwards. Since the GABA_A_ reversal potential in newborn young GCs is depolarized until 3 wpm relative to resting potential (E_GABA_ = -35 mV vs V_m_ = - 80 mV), GABAergic synaptic responses are depolarizing. In the dentate gyrus, PV-basket cells are easily recruited in a feedforward manner during afferent cortical activity (Ewell and Jones, 2010; Groisman et al., 2020). Our data show that a small number of active PV-interneuron synapses (∼3-5) will support the activation of NMDA receptors and thereby promote synaptic integration and cell survival of young GCs during the first 2-3 weeks post mitosis (Tashiro et al., 2006, 2007; Alvarez et al., 2016). In addition, depolarizing GABAergic potentials are expected to promote glutamatergic synapse unsilencing and NMDAR-dependent long-term potentiation (Ge et al., 2007; Kheirbek et al., 2012; Chancey et al., 2013; Li et al., 2017). Thereby, the GABAergic boosting of NMDAR activation generates an advantage for young GCs for activity-dependent glutamatergic synapse formation in comparison to fully mature GCs (Tashiro et al., 2006; Toni et al., 2007; Adlaf et al., 2017).

In case of increased hippocampal activity, additional recruitment of feedback inhibition via PV- and SOM-interneurons is expected (Pernía-Andrade and Jonas, 2014; Temprana et al., 2015; Espinoza et al., 2018; Groisman et al., 2020). With more than about 10-15 active GABAergic synapses, lateral feedback inhibition will effectively shut down AP firing via shunting inhibition in spite of depolarizing GABA (Fig. 7). We have identified one specifically important property of the α5-GABA_A_Rs contributing to this phenomenon, which is the 4-fold amplification of conductance at depolarized potentials close to the AP threshold (V_thresh_= - 35mV, Heigele et al., 2016). As a consequence, GABAergic boosting of AP firing in young GCs is limited to low interneuron activity, whereas shunting inhibition dominates during enhanced hippocampal network activity. This mechanism should reliably ensure and maintain sparse firing of the young GC population under varying conditions *in vivo*.

At 4 wpm, the GABA reversal potential in young GCs is shifted towards the resting potential similar to fully mature neurons (Ge et al., 2006; Heigele et al., 2016). Interestingly, this shift in E_GABA_ is paralleled by the exclusion α5-GABA_A_Rs from perisomatic PV-interneuron synapses (this study) and an increase and speed up of PV-interneuron mediated synaptic transmission (Groisman et al., 2020). Single channel analysis in rat hippocampal brain slices revealed two different populations of GABA_A_ receptors at GC somata: one with linear and one with a voltage-dependent outward rectifying conductance (Birnir et al., 1994). Our new results indicate that these two channel populations could represent synaptic and extrasynaptic GABA_A_Rs at the mature GC soma. It is not fully clear under which physiological conditions extrasynaptic GABA_A_Rs are activated in GCs. However, it was shown that a brief high-frequency burst in dentate gyrus basket cells can generate spillover in basket cell-GC synapses (Overstreet and Westbrook, 2003) and could thus potentially activate extrasynaptic α5-GABA_A_Rs also in mature GCs during particularly strong hippocampal network activity.

In contrast to the pronounced developmental changes in perisomatic synapses, all GCs continue to incorporate α5-GABA_A_Rs in dendritic SOM-interneuron synapses. Our modelling results show that the nonlinear voltage-dependence of these receptors is also beneficial for sparse AP firing in mature cells - although the effect is less pronounced than in younger neurons with depolarized E_GABA_. Interestingly, knockout of α5-GABA_A_Rs selectively in dentate gyrus GCs was shown to double the number of c-fos-positive GCs during exploration of a novel environment (Engin et al., 2015). This was attributed to reduced expression and activation of extrasynaptic GABA_A_Rs. However, our data show that α5-GABA_A_Rs mediate synaptic inhibition by dendrite-targeting SOM interneurons. The same interneurons have been shown to powerfully control the size of active GC cell assemblies (Stefanelli et al., 2016). Therefore, the present results suggest that the nonlinear α5-GABA_A_Rs in SOM-interneuron synapses may be more relevant than extrasynaptic receptors for the control of sparse AP firing in hippocampal GCs.

What is the role of sparse AP firing in young and mature GCs for learning and memory? Behavioral studies showed that newly generated GCs improve the unequivocal distinction of similar memory items, a process called pattern separation (Clelland et al., 2009; Creer et al., 2010; Sahay et al., 2011). Furthermore, GCs have been shown to contribute to pattern separation preferentially when they are young at about 3-4 weeks post mitosis (Gu et al., 2012; Nakashiba et al., 2012). On the network level, sparse and orthogonal firing in hippocampal GCs was shown to improve hippocampal pattern separation (Treves and Rolls, 1992, 1994). This suggests that newborn GCs should be sparsely active during hippocampus-dependent learning in order to support pattern separation.

Studies using immediate early gene expression as a proxy for spiking activity *in vivo* indeed indicate that young GCs are sparsely active. These studies showed that the proportion of active 4-week-old GCs is slightly larger (∼4%) than the proportion of active mature GCs under identical conditions (∼1-2%)(Kee et al., 2007; Krzisch et al., 2016). Similarly, *in vivo* calcium imaging has shown that cells younger than 6 wpm are on average 1.5-fold more active than mature GCs (Danielson et al., 2016). Although slightly more active than mature GCs, the activity levels in young GCs are still one order of magnitude lower than reported activities in CA1 or CA3 (30-40%, Chawla et al., 2005; Leutgeb et al., 2007; Hainmueller and Bartos, 2018). Finally, the sparse and orthogonal AP firing in young GCs is required for beneficial effects of adult neurogenesis on learning. Increasing the active proportion of young cells during learning from 4% to about 9% Arc^+^ neurons via Neuroligin-2 overexpression impairs hippocampus-dependent memory, rather than improving it (Krzisch et al., 2016). Similarly, sparse activity of 4 wpm GCs is necessary for memory consolidation during REM sleep, while optogenetically enhancing the activity in these cells disturbs consolidation (Kumar et al., 2020 in press). This suggests that α5-GABA_A_R-dependent sparsification of young GC firing may be critically important for the beneficial contribution of young GCs to improved pattern separation.

Taken together, our data explain how depolarizing GABAergic synapses can generate a fine-tuned excitation-inhibition balance in immature neurons to enable trophic GABAergic depolarization but avoid runaway excitation, hyperexcitability and epilepsy. We have identified synaptic α5-GABA_A_Rs as important elements to ensure sparse synaptic recruitment of the young hippocampal GCs and show that the nonlinear gating of theses receptors is a crucial step for inhibition of AP firing with depolarized E_GABA_.

## Acknowledgements

We would like to thank Selma Becherer for mouse genotyping, histochemical stainings and excellent technical assistance. This work was supported by the Swiss National Science Foundation (SNSF, Project 31003A_176321). M-C.H is an employee at F. Hoffmann-La Roche. M.L., J.M.S. and J.B. declare no competing financial interests.

## Author contributions

Acquisition and analysis of experimental data: M.L.. Computational modelling: J.M.S.. Study concept and design: J.M.S. and J.B.. Interpretation of results as well as drafting and critical revision of the manuscript: all authors.

## Materials and Methods

### Animals

Experiments were performed on transgenic mice of both genders, expressing the red fluorescent protein DsRed under the control of the doublecortin (DCX) promoter (Couillard-Despres et al., 2006), allowing for visual identification of adult-born granule cells in acute brain slices. Optogenetic experiments were performed with virally injected animals (PVCre x DCX-DsRed or SOMCre x DCX-DsRed). To generate lines for viral injection, homozygous PV-Cre (B6;129P2-*Pvalb*^*tm1(Cre)Arbr*^/J) or SOM-Cre (*SST* ^*tm2.1(cre)Zjh*^/J) mice were crossed with homozygous DCX-DsRed mice (*Dcx-DsRed*, Couillard-Despres et al., 2006). All mice were obtained from The Jackson Laboratory, except for the DCX-DsRed animals, which were generated by Ludwig Aigner (PMU, Salzburg). In order to increase adult neurogenesis, mice were housed in an enriched environment for a minimum of 10 days prior to experiments with running wheels, tunnels and houses. Groups of 3-6 animals were placed in large cages (595×380×200 mm) on a 12:12hr light/dark cycle. Animals had ad libitum access to food and water. All experiments were approved by the Basel Cantonal Committee on Animal Experimentation according to federal and cantonal regulations.

### Stereotaxic viral injections

To allow for selective excitation of PV or SOM interneurons, a floxed-channelrhodopsin (AAV-EF1a-DIO-hChR2(H134R)-EYFP, UNC Vector Core) was expressed in the dentate gyrus of PV-Cre x DCX-DsRed or SOM-Cre x DCX-DsRed mice. Four- to five-week-old mice were anaesthetized using inhalation of 4% isoflurane (2% for maintenance) and placed in a stereotaxic frame to level the skull using bregma and lambda. A small incision was made in the skin over the skull and 0.8 mm holes were drilled above the injection site. Injections were made using pulled glass micropipettes (BLAUBRAND Intramark micropipettes) and a picospritzer (version III Parker Hannifin, 2-15 ms air pulses at 1Hz, between 10-20 psi). The following co-ordinates for the ventral dentate gyrus were used, with four injection sites along the dorso-ventral axis: anterior-posterior= -3.2mm, medial-lateral= ±3.0mm, dorsal/ventral=-4.5mm, -4.0mm, -3.5mm and -3.00mm.

### Slice preparation for patch-clamp recordings

Adult 5- to-10-week-old male and female mice were anaesthetized with isoflurane (4% in O2, Vapor, Draeger) and once sufficiently unresponsive, killed by decapitation in accordance with national and institutional guidelines. To increase cell survival, animals were placed in an oxygen-enriched environment for a minimum of ten minutes prior to anesthesia. Horizontal brain slices were prepared as previously described (Bischofberger et al., 2006). Briefly, the brain was removed in an ice-cold sucrose-based solution (87 NaCl, 25 NaHCO_3_, 2.5 KCl, 1.25 NaH_2_PO_4_, 75 sucrose, 0.5 CaCl_2_, 7 MgCl_2_ and 10 glucose, Osmolarity 325-328 mOsm approximately 4°C), which was aerated with carbogen (a mixture of 95% O_2_ and 5% CO_2_). Transverse 350μm thick hippocampal slices were cut using a Leica VT1200 vibratome with a cutting velocity of 0.04 mm/s and a horizontal vibration amplitude of 1.8 mm (vertical vibrations < 1 µm). Slices were incubated at 35°C in the same sucrose-based solution for 30 minutes and then stored at room temperature until experiments were conducted.

### Patch-clamp recordings

Electrophysiological recordings of acute brain slices were conducted within about 6 hours after slice preparation. Slices were placed into a bath chamber, with continuous perfusion of oxygenated artificial cerebrospinal fluid. The ACSF contained (in mM) 125 NaCl, 25 NaHCO_3_, 25 glucose, 2.5 KCl, 1.25 NaH_2_PO_4_, 2 CaCl_2_, and 1 MgCl2 (equilibrated with 95% O_2_/5% CO_2_, Osmolarity 315-25 mOsm). Granule cells were visualized with an infrared differential interference contrast (IR-DIC) using a 63x objective (numerical aperture 1.0, Zeiss, Oberkochen, Germany). Adult-born granule cells between 1 and 4 weeks of age were identified by detection of DsRed fluorescence using a cooled CCD camera system (SensiCam, TILL Photonics, Gräfelfing, Germany). DsRed positive cells were detected using a light source with an excitation wavelength of 555nm (Polychrome V, TILL Photonics), which was connected to the microscope using a quartz fiber optic light guide. The illumination intensity was kept low to avoid possible photo-bleaching of the neurons and subsequent phototoxicity. Cell maturity and age of DsRed expressing young cells was further constrained using the electrical input resistance. DsRed-positive neurons with an input resistance between 2-8 GΩ were classified as 2-3 weeks old, while 0.5-2 GΩ cells were classified as 3-4 weeks old. Mature cells lacked DsRed fluorescence and had an input resistance below 400 MΩ (Schmidt-Hieber et al., 2004; Heigele et al., 2016; Li et al., 2017).

Voltage-clamp recordings of young and mature granule cells were conducted with patch pipettes (3-6 MΩ) pulled from borosilicate glass tubes with 2.0 mm outer diameter and 0.5 mm wall thickness (Hilgenberg, Malsfeld, Germany; Flaming-Brown P-97 puller, Sutter Instruments, Novato, USA). Current-clamp recordings of newborn granule cells were conducted using glass with 0.7 mm wall thickness with higher pipette resistances (6-12MΩ) and low capacitance (≈5pF).

Voltage-clamp recordings for connectivity and rectification experiments of both young and mature granule cells were conducted with high chloride internal solutions to increase the amplitudes of inward currents recorded from young granule cells. A potassium-chloride based intracellular solution containing symmetrical chloride was used resulting in a reversal potential of about 0 mV (in mM: 140 KCl, 10 EGTA, 10 HEPEs, 2 MgCl_2_, 2 Na_2_ATP, 1 Phosphocreatine, 0.3 GTP, adjusted to pH 7.3 with KOH). To determine the physiological level of rectification in young cells, we used an internal solution with physiological levels of chloride (in mM: 122 K-Gluconate, 21 KCl, 10 HEPEs, 10 EGTA, 2 MgCl_2_, 2 Na_2_ATP, 1 Phosphocreatine, 0.3 GTP adjusted to pH 7.3 with KOH) in order to obtain a GABA reversal potential comparable to perforated-patch measurements (Heigele et al., 2016). Additionally, we used this internal solution for all current clamp recordings. For voltage-clamp recordings targeting dendritic inputs, cesium-based internal solutions were used to block dendritic potassium channels and increase voltage control during dendritic GABA_A_ currents. For rectification experiments pipettes were filled with (in mM): 100 CsCl, 40 Cs-gluconate, 10 HEPEs, 10 EGTA, 2 MgCl_2_, 2 Na_2_ATP, 1 Phosphocreatine, 0.3 GTP, adjusted to pH 7.3 with CsOH. For pharmacological experiments of dendritic inputs onto mature GCs pipettes were filled with (in mM): 135 Cs-gluconate, 2 CsCl, 10 HEPEs, 10 EGTA, 2 MgCl_2_, 2 Na_2_ATP, 2 TEACl, adjusted to pH 7.3 with CsOH. In some internal solutions we added 5 mM QX314-Cl to block voltage-gated sodium channels and prevent action currents in voltage clamp. In this case, low-chloride solutions were adjusted to maintain the original chloride concentration.

GABA-mediated synaptic transmission was isolated by pharmacological block of glutamatergic currents using 10 μM NBQX (2,3-dioxo-6-nitro-1,2,3,4-tetrahydrobenzo[*f*]quinoxaline-7-sulfonamide) and 25 μM AP5 (D-(−)-2-amino-5-phosphonopentanoic acid). Gabazine (SR95531, 0.2–10 μM) was used to block GABA_A_-mediated currents as indicated.

Voltage and current clamp recordings were both obtained using a Multiclamp 700B amplifier (Molecular devices Devices, Palo Alto, CA, USA), filtered at 10 kHz, and digitized at 20kHz with a CED Power 1401 interface (Cambridge Electronic Design, Cambridge, UK). Bridge balance was used to compensate the series resistance (R_s_) in current clamp recordings (R_s_ ≈ 10-50 MΩ). In most voltage-clamp recordings, series resistance was compensated with an 80% correction. All voltage-clamp experiments were performed at room temperature (22-24°C), while current clamp recordings were obtained at high temperature (30-33°C). Data acquisition was controlled using IGOR Pro 6.31 (WaveMetrics, Lake Oswego, Oregon) and the CFS library support from CED (Cambridge Electronic Design, Cambridge, UK).

### Extracellular synaptic stimulation

To electrically stimulate synaptic inputs, 12-15 MΩ pipettes were filled with a HEPES-buffered sodium rich solution to apply short negative current pulses (5-60 μA, 200 μs). To stimulate axons of soma-targeting GABAergic interneurons, the stimulation pipette was placed in the outer third of the granule cell layer (GCL), with a minimal lateral distance of 100 μm from the recorded cell. To stimulate axons of dendrite-targeting GABAergic interneurons, the stimulation pipette was placed in the outer third of the molecular layer (ML). For current clamp experiments, stimulation of perforant path glutamatergic fibers and dendrite-targeting GABAergic fibers was done by placing the stimulation pipette in the middle of the ML.

### Channelrhodopsin-mediated activation of GABAergic interneurons

A diode laser (DL-473, Rapp Optoelectronic) was coupled to the epifluorescent port of the microscope (Zeiss Examiner, equipped with a 63x NA1.0 water immersion objective; Carl Zeiss Microscopy GmbH, Jena, Germany) via fiber optics. The laser was controlled via TTL pulses. The field of illumination was targeted to the granule cell layer when stimulating parvalbumin interneurons, and the hilus or outer molecular layer when targeting somatostatin positive interneurons. For rectification experiments the intensity of the light-pulse was adjusted to produce a post-synaptic current of around 200 pA at -80mV (5-20mW, 5ms).

### Drugs and reagents

D-AP5 (50 mM; Tocris) and SR95531 (0.2μM and 10μM, Tocris) were dissolved in water. CGP54626 hydrochloride (10 mM; Tocris), NBQX (20 mM; Tocris), and the α5-NAM RO4938581 (10 mM; F. Hoffmann-La Roche) were dissolved in DMSO. QX314 (5mM, Tocris) was dissolved in the internal pipette solution. All other chemicals were obtained from Sigma or Merck.

### Immunohistochemistry

Immunohistochemical analysis was done with 350μm horizontal slices taken during electrophysiological preparation. Slices were fixed in 4% paraformaldehyde for 90 min to maintain the integrity of the postsynaptic density. Washing was done with a step-wise protocol using a tris-buffered saline, and 1% triton solution. Slices were transferred to the same tris-buffered saline containing Bovine serum albumin (BSA, 1%) for two hours to block unspecific binding of antibodies. Incubation with a primary antibody (guinea pig-anti-VGAT, 1:500; rabbit-anti-GABA_A_ receptor α5, 1:500 both from Synaptic Systems) in 1% BSA was done for 24-48 hours at 4°C. Subsequently, slices were rinsed with tris-buffered saline and incubated with the secondary antibodies at room temperature for 2 hours, including donkey-anti-rabbit-Alexa Fluor 488 (1:1000, MoBiTec), donkey-anti-guinea pig Alexa Fluor 647 (1:1000, Millipore Bioscience Research Reagents), Alexa Fluor 568 conjugated Streptavidin (MoBiTec), and DAPI (1:10,000, Sigma Aldrich). After the final rinsing, slices were mounted with ProLong Gold Antifade (Invitrogen), and imaged using a Zeiss LSM700 confocal microscope (Oberkochen, Germany). Image analysis and processing was done using the Zeiss ZEN software and ImageJ freeware (https://imagej.net/).

### Data analysis and statistics

Analysis of patch-clamp data was performed using the open source analysis software Stimfit (Guzman et al., 2014, http://code.google.com/p/stimfit) and customized scripts written in Python.

#### Intrinsic cell properties

The intrinsic properties of recorded neurons were determined in the first initial minutes after establishing the whole-cell configuration. Once the membrane was ruptured, the input resistance was measured in voltage-clamp using the current response to a negative voltage step (−5 mV, 500 ms pulse) from a holding potential of -80 mV. Action potential properties were determined in current-clamp using 500-ms-current pulses (4-50pA) to elicit a single AP.

#### Analysis of synaptic currents

For IPSCs, the decay τ was calculated as the amplitude-weighted average of a biexponential fit to the decay phase of the PSC starting at 95% of its amplitude. To visualize the voltage-dependent rectification, the synaptic currents were normalized by the maximal outward current and fitted to the following sigmoidal function using GraphPad Prism: 

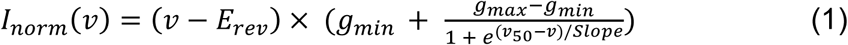

with *v* representing the membrane potential, E_rev_ the GABA_A_ reversal potential, g_min_ and g_max_ the minimal and maximal conductance, *v*_50_ the membrane potential at half-maximal conductance increase, and Slope determining the steepness of the sigmoidal, which was constrained to be at least the voltage difference between two adjacent data points.

To calculate the rectification index (RI) for each individual cell, we first calculated the maximal slope conductance obtained from a linear fit to the outward currents (V_m_ > 0 mV). This conductance was used to extrapolate a predicted current amplitude at -80 mV, which was divided by the actually measured value at –80 mV (1 = no deviation from linear).

#### PSP analysis

Mean burst PSPs were analyzed to obtain peak amplitude and integral. The analysis of burst PSPs in current-clamp recordings testing shunting inhibition was performed on single-trial data to avoid distortion by AP discharge. In cases of AP firing, APs were digitally removed by cutting off spikes at the AP threshold, defined by the voltage slope (10 V/s) before calculating the integral.

#### Statistical analysis

Statistical analysis was performed with GraphPad Prism 8. We did not rely on the assumption that our data followed a normal distribution, and thus used non-parametric tests. In most instances, statistical significance of paired data, in particular normalized data relative to 100% control, was derived from the Wilcoxon matched-pairs signed-rank test. The Mann-Whitney test was used for all unpaired comparisons. A two-tailed test was used. The significance level was set to P=0.05. All data are shown as mean ± s.e.m if not stated otherwise. The sample size was determined by the reproducibility of the experiments and based on our experience with similar experiments (Heigele et al., 2016; Li et al., 2017; Schulz et al., 2018). The number *n* of observations reflects the number of cells recorded from.

### Computational modelling

We designed two GC models in the NEURON simulation environment (Hines and Carnevale, 1997). The mature GC model reflecting a representative mature mouse granule cell was generated based on previously developed passive cable models (Schmidt-Hieber et al., 2007; Schmidt-Hieber and Bischofberger, 2010). Accordingly, the specific membrane capacitance was set to 1 μF/cm_2_, the specific membrane resistivity to R_m_ = 40 kΩcm^2^, and the specific internal resistivity to R_i_ = 200 Ωcm. To indirectly model the effect of synaptic spines on membrane properties the membrane capacitance in the dendrites was increased by a factor of 1.5. Similarly, the specific membrane resistivity was decreased by the same factor. A model for a prototypical adult born 2.5 week-old was constrained by previously obtained experimental data on passive an active membrane properties of young GCs (Heigele et al., 2016). The cell’s capacitance is closely correlated to R_in_ via the age-dependent membrane time constant τ_m_ (Heigele et al., 2016) and largely reflects the total membrane area. The specific membrane capacitance was assumed to be identical to mature GCs (1 μF/cm^2^) and the specific membrane resistivity was deduced from the age-dependent τ_m_. Using this set of parameters, the dimensions of the morphology were constrained for a fixed branching pattern. The young GC model (about 2.5 wpm) had a representative R_in_ of 4 GΩ while the model of mature GC showed a R_in_ of about 200 MΩ. The resting membrane potential for young and mature cells was set to E_Leak_ = -80 mV (Heigele et al., 2016).

Glutamate synapses, including an AMPAR- and a NMDAR-mediated conductance, and GABA synapses were distributed in a fixed pattern over the dendritic tree depending on stimulation paradigm. AMPA, NMDA and GABA conductances were modeled with exponential rise and decay using the Exp2Syn class. For the AMPA conductance, the time constant (τ) for rise was set at 0.2 ms and the decay τ at 2 ms; for the NMDA conductance the rise τ was set to 3 ms and the decay τ to 35 ms (Schulz et al., 2018). The voltage-dependent magnesium (Mg^2+^) block of NMDARs was modeled as

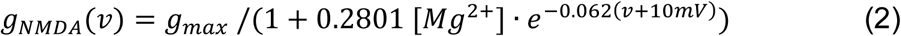

(Jahr and Stevens, 1990; Schulz et al., 2018). The extracellular magnesium concentration [Mg^2+^] was set to 1 mM according to the concentration in our ACSF.

Nonlinear GABAergic synapses that showed outward rectification as observed experimentally (Fig.1) were modelled as the sum of two Exp2Syn class synapses in NEURON as published previously (Schulz et al., 2018). Briefly, the first component was modeled as a linear conductance and was assigned 20% of the total synaptic weight. The second rectifying conductance component was assigned 80% of the total synaptic weight. The 4-fold voltage-dependent outward rectification of this second component was modeled as

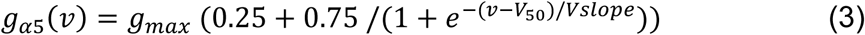

Based on the fit to the experimentally determined voltage-dependence of pharmacologically isolated α5-GABA_A_Rs-mediated IPSCs (Schulz et al., 2019), *V*_*slope*_ was set to 7 mV and *V*_*50*_ to -45 mV. For the linear GABA synapse, rise and decay τ were set to 0.5 and 10 ms, respectively; for the outward rectifying GABA synapse, the rise and decay τ were set to 3 and 55 ms, respectively.

The maximal conductance of synaptic receptors in the mature GC model was set to the same values as in our previously published CA1 pyramidal neuron model (Schulz et al., 2018). For young GCs, the AMPAR conductance was decreased from 0.14 nS to 0.02 nS to account for the very small EPSCs and the large NMDA/AMPA ratio observed in young GCs (Li et al., 2017). The GABA synapse conductance in the young GC model was adjusted from 0.7 nS to 0.24 nS, so that the activation of individual GABA synapses evoked a PSP of about 3.3 mV similar to spontaneous GABA-mediated PSPs (3.31±0.21 mV, n=5) in the presence of glutamate receptor blockers during current-clamp recordings with physiological chloride concentration. To explore the functional impact of the voltage-dependent nonlinearities of GABAergic synapses, the synapses were linearized by keeping the nonlinear component constant to the small conductance corresponding to -80 mV (i.e. 0.28 and 0.096 nS in mature and young GC model, respectively).

To study the effect of GABA-mediated inhibition on AP output (Fig. 7), we included voltage-dependent sodium (Na_V_) and delayed rectifier potassium channels (K_DR_). The eight state Na_V_ model reported in Schmidt-Hieber & Bischofberger (2010) was included in two versions, to account for the functional properties of somato-dendritic and axonal sodium channels in mature mouse dentate gyrus granule cells. All gating parameters were taken directly from Schmidt-Hieber & Bischofberger (2010). A kinetic scaling factor was added to enable a slowing of the gating kinetics in young GCs by a factor of 2 to account for the much slower AP waveforms. Activation and inactivation of Na_V_ were shifted by 12 mV and 26.5 mV (+12 mV due to Donnan potential) to the right in the mature and young GC model, respectively, to ensure AP firing at the physiologically defined current (mature, 113 pA; young, 8.5 pA) and voltage thresholds (mature, -50 mV; young, -35 mV) during 1 sec long current steps (Heigele et al., 2016). Persistent and transient K_DR_ were taken from Hay et al. (2013) and included at a fixed ratio 100:1 of persistent versus transient K_DR_. Sodium and potassium reversal potentials were set to E_Na_ = 60 mV and E_K_ = -90 mV, respectively. The axon initial segment (AIS) contained the highest density of voltage-dependent conductances. In the other compartments the channel density was reduced according to previously published ratios (Schmidt-Hieber & Bischofberger 2010). Table 1 summarizes the channel densities used.

**Table 1.**
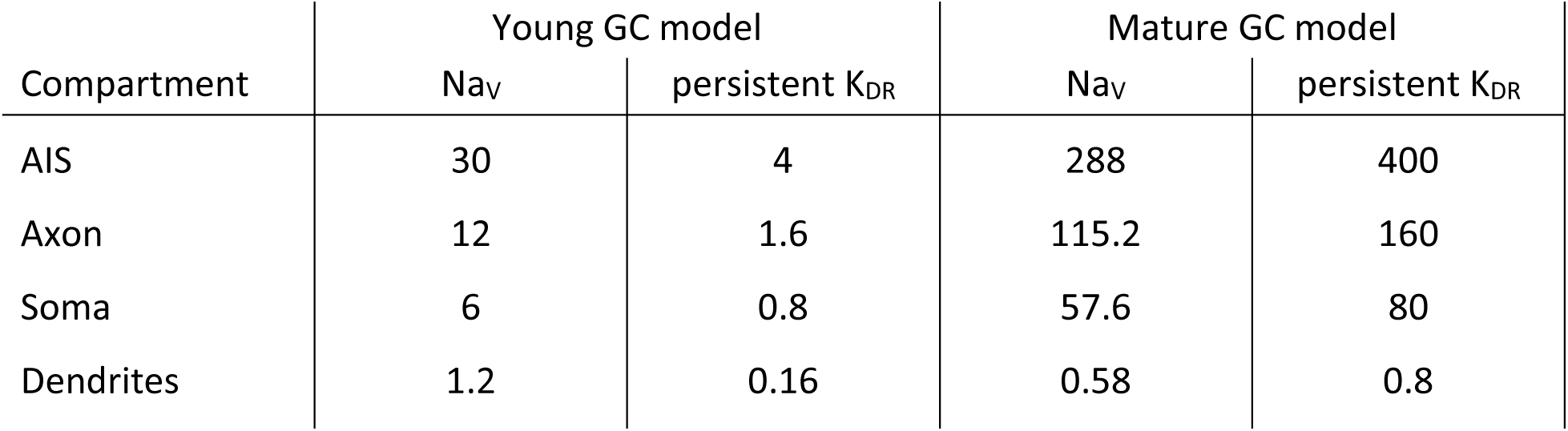
Density (mS/cm^2^) of Na_V_ and persistent K_DR_ channels in the different compartments of the young and mature GC models.

| Compartment | Young GC model |  | Mature GC model |  |
| --- | --- | --- | --- | --- |
| | $Na_V$ | persistent $K_{DR}$ | $Na_V$ | persistent $K_{DR}$ |
| AIS | 30 | 4 | 288 | 400 |
| Axon | 12 | 1.6 | 115.2 | 160 |
| Soma | 6 | 0.8 | 57.6 | 80 |
| Dendrites | 1.2 | 0.16 | 0.58 | 0.8 |

## Legends

**Supplementary Fig. 1.**
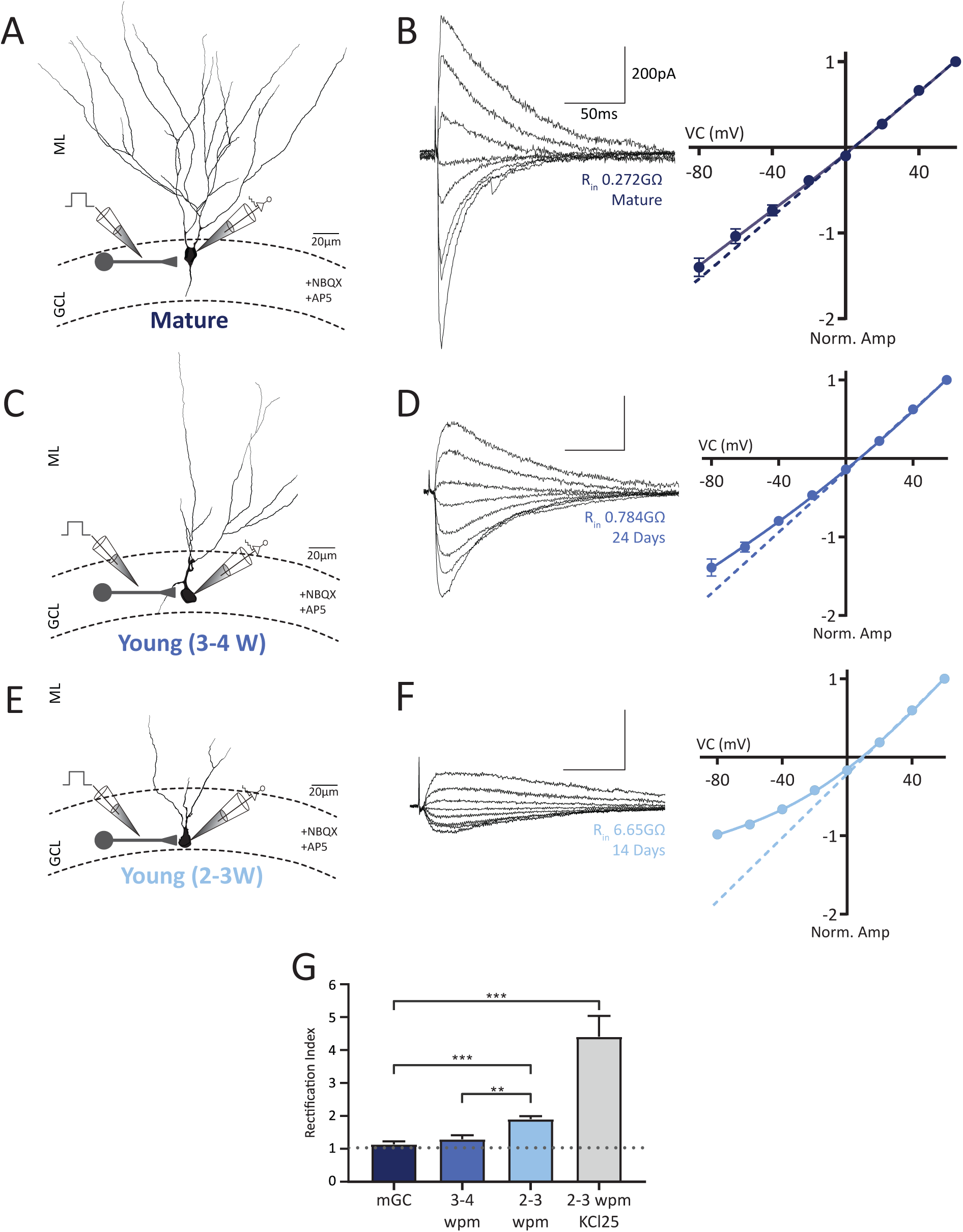
Young but not mature GCs receive non-linear outward rectifying GABAergic inputs activated by GCL stimulation. **A, C, E**) Experimental design. Electrical stimulation of the granule cells layer (GCL) was used to target somatic inputs onto mature (A) and young GCs (C, E) (5-20μA). Cells were classified based on DCX-DsRed fluorescence and the measured input resistance. Mature cells were defined as having an input resistance less than 400MΩ, young cells with an input resistance between 500MΩ and 2GΩ correspond to 3-4 weeks of age, and young GCs with input resistances between 2 and 8 GΩ are corresponding to 2-3 weeks of age. All young cells recorded were also DCX-DsRed positive. **B**) GPSCs from mature GCs were recorded at increasing membrane potentials (−80mV to +60mV) in symmetrical chloride conditions. The dashed line represents a linear fit to the outward current (+20 to +60mV), while the solid line represents a non-linear fit obtained with the equation (1) outlined in methods. Recordings were done in the presence 10 µM NBQX and 25 µM AP5. **D, F**) as in B, but for young GCs. The IV curve of synaptic current in young GCs clearly shows the non-linearity of the recorded GPSCs. **G**) Mean rectification indices (RI) were quantified as the expected conductance extrapolated from the linear fit to the outward currents (above 0mV) divided by the measured value at -80mV (where 1= no deviation, indicated by grey dashed line). The mean RI was significantly larger in young GCS than in mature.

## Notes

### Competing Interest Statement

The authors have declared no competing interest.

